# Disruption of Reelin signaling in a dual-hit mouse model of schizophrenia: impact of postnatal Δ9-tetrahydrocannabinol exposure in a maternal immune activation model

**DOI:** 10.1101/2025.07.25.666721

**Authors:** Celia Martín-Cuevas, Víctor Darío Ramos-Herrero, Álvaro Flores-Martínez, Irene González-Recio, María Luz Martínez-Chantar, Juan Carlos Leza, J. Javier Meana, Benedicto Crespo-Facorro, Ana C. Sánchez-Hidalgo

**Affiliations:** Instituto de Biomedicina de Sevilla (IBiS)/University Hospital Virgen del Rocío/CSIC/University of Sevilla, Manuel Siurot AV, 41013, Seville, Spain; Spanish Network for Research in Mental Health (CIBERSAM, ISCIII), Monforte de Lemos AV, 3-5, 28029, Madrid, Spain; Department of Psychiatry, School of Medicine, University of Sevilla, Manuel Siurot AV, 41013, Seville, Spain; Liver Disease Laboratory, Center for Cooperative Research in Biosciences (CIC bioGUNE), Basque Research and Technology Alliance (BRTA), Derio (Bizkaia), Spain; Center for Biomedical Research in Liver and Digestive Diseases Network (CIBERehd), Bizkaia, Spain; Department of Pharmacology, Faculty of Medicine, Universidad Complutense, Madrid 28040, Spain; Department of Pharmacology, University of the Basque Country UPV/EHU, Leioa, 48940, Bizkaia, Spain; Biobizkaia Health Research Institute, Barakaldo, 48903, Bizkaia, Spain

**Keywords:** poly(I:C), cannabis, behavior, brain, proteomics, neurodevelopment, extracellular matrix, psychosis

## Abstract

Since the discovery of the first antipsychotic, pharmacological treatment of schizophrenia (SCZ) has primarily relied on agents that block D2 dopamine receptors. However, due to variability in patient responses, there is a pressing need to identify new biomarkers and therapeutic strategies. In this context, we have developed a dual-hit mouse model that combines maternal immune activation (MIA) induced by polyinosinic-polycytidylic acid (Poly(I:C)) and postnatal exposure to Δ9-tetrahydrocannabinol (THC), the psychoactive component of cannabis. We assessed the face validity of this model and investigated potential alterations in the Reelin signaling pathway. Our findings show a reduction in Reelin levels, a new potential key biomarker of SCZ, in the prefrontal cortex of male mice treated with THC compared to the dual-hit group, and across all treatment groups compared to controls in females. Additionally, a decrease in the number of Reelin+ cells was observed across these groups. The dual-hit model exhibited phenotypes indicative of positive symptoms in males, as well as phenotypes associated with negative symptoms in both sexes. Furthermore, the model demonstrated reduced cortical thickness in THC-treated groups, alongside decreased dendritic spine density in both the prefrontal cortex and hippocampus in the dual-hit group.

## 1. Introduction

Since the discovery of the first antipsychotic drug, chlorpromazine, in 1952, pharmacological treatment of schizophrenia (SCZ) has predominantly relied on agents that block D2 dopamine receptors (Carlsson & Carlsson, 2006). However, patient responses remain highly variable, with only about 60% achieving significant clinical improvement, while approximately 30% remain resistant to standard treatments (Crespo-Facorro et al., 2007; Moreno-Sancho et al., 2022). The disorder exhibits a higher incidence in men, who tend to present earlier onset and greater severity (Aleman et al., 2003; Eranti et al., 2013; Mendrek & Mancini-Marïe, 2016; Shangase et al., 2023), suggesting a clear sexual dimorphism. This variability in treatment response underscores the critical need for a more nuanced understanding of the biological mechanisms underlying SCZ to facilitate the identification of more targeted and effective therapeutic approaches.

SCZ is a chronic, severe psychiatric disorder characterized by a heterogeneous combination of symptoms (Patel et al., 2014; Haddad & Correll, 2018; American Psychiatric Association, 2021), typically categorized into positive, negative, and cognitive (Rodríguez-Sánchez et al., 2007; Patel et al., 2014; McCutcheon et al., 2020; Crespo-Facorro et al., 2021). It is caused by both genetic and environmental factors, particularly during the prenatal stages (Patel et al., 2020; American Psychiatric Association, 2021; Martín-Cuevas et al., 2023). These findings have led to the neurodevelopmental hypothesis of the disorder, which postulates that prenatal and perinatal disturbances, whether genetic or environmental, interact with other environmental factors to trigger the onset of the disease at some point during development (Guerrin et al., 2021).

The study of SCZ through animal models allows for a faster and more controlled analysis of the structural and molecular alterations associated with the disorder, as well as the testing of behavioral phenotypes and new therapeutic molecules (MacDowell et al., 2017; Shangase et al., 2023). Most current models include only one type of alteration (either genetic or environmental) and fail to reproduce the full spectrum of symptoms seen in the disorder. Therefore, combining multiple risk factors may more accurately replicate the clinical presentation of the disease and further validate the hypothesis of neurodevelopmental disturbances. In light of this, the dual-hit hypothesis of SCZ has been proposed (Bayer et al., 1999; Maynard et al., 2001; Suárez-Pinilla et al., 2014; Setién-Suero et al., 2020; Hall et al., 2023). This hypothesis suggests that a prenatal insult, such as an immune challenge during pregnancy leading to maternal immune activation (MIA), predisposes the individual to the development of SCZ (Guma et al., 2019; Bauman et al., 2020; Han et al., 2021a, 2021b), and a second postnatal insult further increases the risk, exceeding the threshold for the onset of the disorder (Malaspina, 1999; Lecca et al., 2019; Martín-Cuevas et al., 2023; Guma et al., 2023; Murlanova & Pletnikov, 2023).

We have established a dual-hit mouse model to study the face validity of the model and to analyze the possible alteration of the Reelin signaling pathway. The model consisted of an induced MIA induced via the injection of polyinosinic-polycytidylic acid (Poly(I:C)), a synthetic double-stranded RNA compound that triggers a signaling cascade leading to the expression of pro-inflammatory cytokines (Haddad et al., 2020; Sánchez-Hidalgo et al., 2022). In the adolescent postnatal stage, the offspring were treated with Δ9-tetrahydrocannabinol (THC), the psychoactive component of cannabis, which acts as an exogenous agonist of the endocannabinoid receptors CB1 (CB1R) and CB2 (CB2R) (Suárez-Pinilla et al., 2014; Lamanna-Rama, et al., 2024). Cannabis use, especially during adolescence, has been shown to increase the risk of developing SCZ (Suárez-Pinilla et al., 2014; Johnson-Ferguson & Di Forti, 2023) with cannabis use during this period resulting in long-term cognitive impairments (Patel et al., 2020), alterations in cortical thickness (Abush et al., 2018), and a reduction in the density and complexity of dendritic spines in the prefrontal cortex (PFC) and hippocampus (HP) (Chesworth & Karl, 2017).

This dual-hit model allows us to study therapeutic targets of SCZ that may facilitate the development of new therapeutic approaches. One potential biomarker of SCZ that has gained increasing attention in recent years is Reelin, a key component of the extracellular matrix (ECM) (Wasser & Herz, 2017; Marzan et al., 2021; Martín-Cuevas et al., 2023). Reelin is a glycoprotein essential for neuronal migration and laminar positioning during brain development (Alexander et al., 2023), regulation of dendritic spine growth and formation, synaptogenesis, synaptic plasticity (Wasser & Herz, 2017; Yamakage et al., 2019; Jossin, 2020), and adult neurogenesis (Bosch et al., 2016).

Reelin binds to apolipoprotein E receptor 2 (ApoER2) and very low-density lipoprotein receptor (VLDLR), which leads to the phosphorylation of the intracellular adaptor Disabled homolog-1 (Dab1). Reelin consists of a signal peptide, an F-spondin homology domain, and a unique region, followed by a main body that includes 8 repeats (R1–R8) and a basic 33-amino acid extension (Bosch et al., 2016; Ogino et al., 2017; Wasser & Herz, 2017; Alexander et al., 2023). Upon secretion, Reelin undergoes processing at two sites, between repeats 2 and 3, and between repeats 6 and 7, yielding five fragments, which are designated N-R2, R3-6, R7- 8, N-R6, and R3-8 (Jossin et al., 2004, 2007, 2020; Hattori & Kohno, 2021). Genetic alterations in Reelin, as well as disruptions in its signaling pathway or reduced expression, have been found in postmortem brains of individuals with SCZ (Howell & Pillai, 2016; Jossin, 2020; Marzan et al., 2021) and Alzheimer’s Disease (AD), where these disruptions correlate with amyloid-beta accumulation and cognitive deficits (Ovadia & Shifman, 2011; Lidón et al., 2020; Valderrama-Mantilla et al., 2025).

In the present study, we focus on investigating novel therapeutic targets to enhance our understanding of SCZ, with a particular emphasis on validating Reelin as a potential biomarker of the disease (Fig. 1).

**Figure 1.**
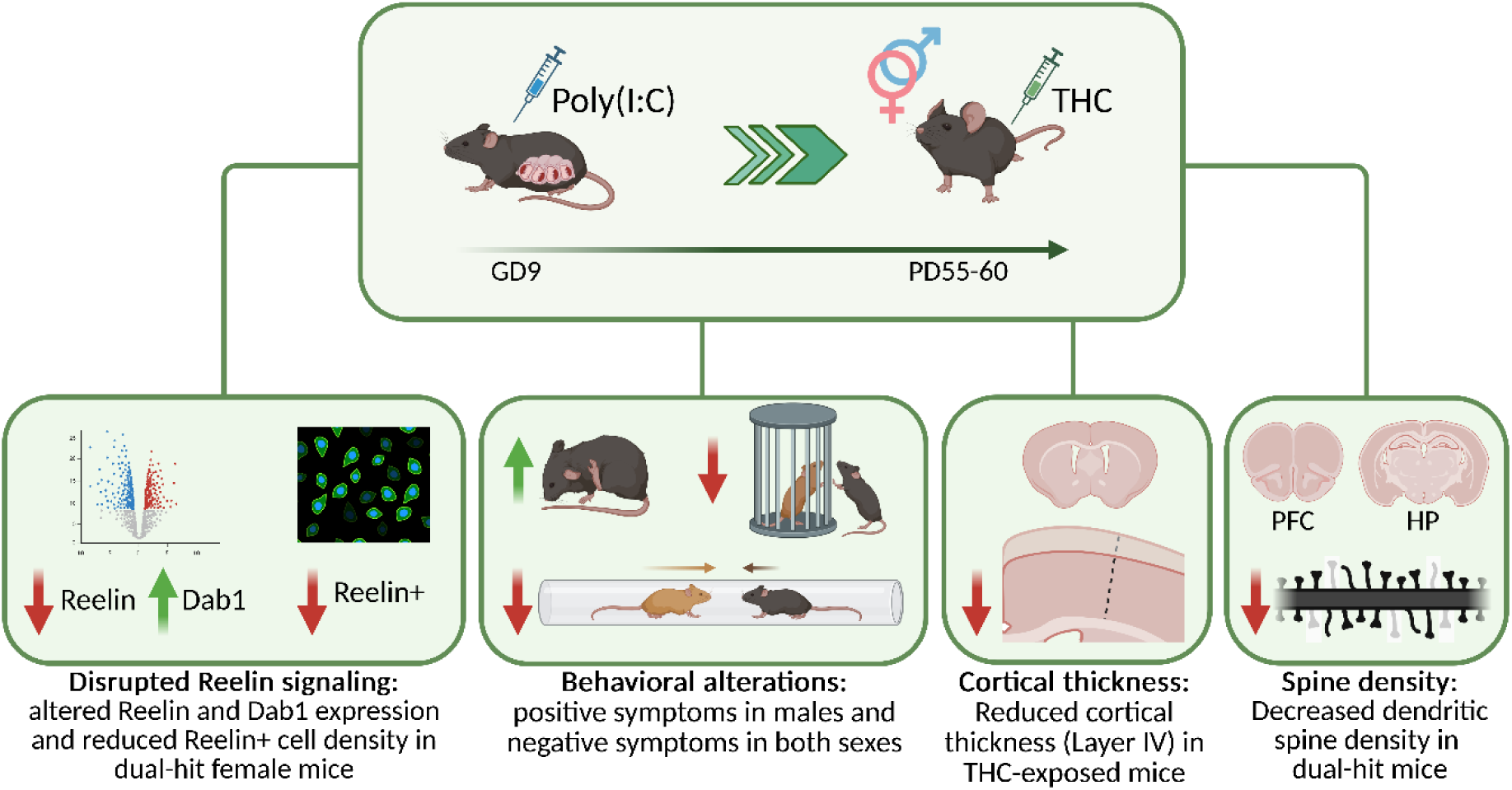
Graphical abstract. Abbreviations: Poly(I:C): polyinosinic-polycytidylic acid; THC: Δ9-tetrahydrocannabinol; GD: gestational day; PD: postnatal day; PFC: prefrontal cortex; HP: hippocampus. Created with Biorender.com

## 2. Materials and methods

### 2.1 Animals

Adult male and female C57BL/6J mice (8-10 weeks old) were purchased from Charles River Laboratories (Saint Germain Muelles, France) and housed in standard laboratory cages in the animal facility of the Instituto de Biomedicina de Sevilla (IBiS), each one containing group of maximum five individuals of the same sex with corncob bedding and environmental enrichment (nesting material and cardboard tunnels). The facility had controlled temperature (22±1°C) and humidity and operated on a 12-hours light/dark cycle (lights on at 8:00 a.m.). Mice had *ad libitum* access to food and water and were acclimated to the facility 1-2 weeks before breeding. For timed mating, one female was placed with one male per cage. Copulatory plugs were checked daily, with GD0 determined by the first detection of a plug. Male mice were separated from females immediately after the defection of a plug to prevent additional mating and reduce potential stress to the pregnant dams. All procedures were conducted in accordance with the ARRIVE and institutional animal care guidelines, and all protocols were approved by the Committee of Animal Use for Research at the Hospital Universitario Virgen del Rocío/Hospital Universitario Virgen Macarena (Spain) (13/10/2021/157).

### 2.2 Dual-hit mouse model paradigm

#### 2.2.1 Maternal Immune Activation

Poly(I:C) sodium salt (P1530-25MG, batch number #0000125513) was purchased from Sigma-Aldrich (Merck KGaA, Darmstadt, Germany) and freshly dissolved daily in sterile saline (0.9% NaCl) at a concentration of 1 mg/ml. On GD9, C57BL/6J pregnant dams were randomly injected intraperitoneally (i.p.) with a single dose of Poly(I:C) (5 mg/kg) or saline (0.9% NaCl). Room temperature at the time of injection was maintained at 22±1°C. To minimize handling stress and its potential confounding effect on the MIA response, pregnant dams received the injection in the same home cages where mating had previously taken place between 9:00 and 10:00 a.m. (Kentner et al., 2018). We selected GD9 for Poly(I:C) injection due to its correspondence with a critical neurodevelopmental period in the mouse cortex, equivalent to the first or second trimester of human pregnancy (Fatemi et al., 2002; Reisinger et al., 2015; Guma et al., 2023) and a dose of 5 mg/kg was chosen based on existing literature (Meyer et al., 2006; Haddad et al., 2020; Guma et al., 2023; Munarriz-Cuezva and Meana, 2025). The same Poly(I:C) batch was used across all experiments to minimize variability in potency between different batches (Mueller et al., 2019).

To assess MIA following Poly(I:C) administration, the weight and temperature of the injected dams were monitored from GD9 to GD11 (pre-injection and at 3, 6, 24 and 48- hours post-injection) (Saline: n = 52; Poly(I:C): n = 53). Clinical signs of sickness behavior (e.g., piloerection, reduced locomotion, huddling) were monitored qualitatively during this period. Two dams per group were sacrificed 3 hours after injection to obtain maternal spleen and embryos (with placenta) (n = 6 mice per treatment) for cytokine level measurements (IL-6 and TNFα) via RT-qPCR. Litters were left undisturbed until PD7, with cage changes performed only one per week thereafter. The offspring were weaned on PD 30 and housed in groups of maximum 5 individuals, separated by sex and litter. Environmental enrichment was maintained throughout the postnatal period. Both male and female were used in subsequent experiments. MIA validation data is available in suppl. Fig. S1. and the list of pregnant dams, injected Poly(I:C) or saline volume and number of offsprings is available in suppl. mat. 2.

#### 2.2.2 THC exposure

THC (T2386) was purchased from Sigma-Aldrich and diluted in vehicle (5% ethanol, 5% cremophor (Sigma-Aldrich), 90% saline). Offspring were assigned to receive daily i.p. injections of THC (10 mg/kg) or vehicle during 6 consecutive days during the adolescent period (PD55-60), modeling subchronic daily use. Four experimental groups were established: Saline + Vehicle (control group), Poly(I:C) + Vehicle, Saline + THC (one-hit groups) and Poly(I:C) + THC (dual-hit group). Adult mice weight was measured on PD55, PD60 and PD75 (n = 12-15 mice per group and sex) (suppl. Fig. S1) and animals were sacrificed on PD75 by cervical dislocation or anesthesia (Tiobarbital, Braun) for transcardiac perfusion.

### 2.3 Behavioral Assays

Behavioral studies were conducted on 107 mice from the four experimental groups previously described (Males: Saline + Vehicle n = 14; Poly(I:C) + Vehicle n = 13; Saline + THC n = 15; Poly(I:C) + THC n = 14. Females: Saline + Vehicle n = 12; Poly(I:C) + Vehicle n = 13; Saline + THC n = 14; Poly(I:C) + THC n = 12) between PD61 and PD74 (one test per day). All behavioral tests were performed between 9 a.m. and 5 p.m. under dim light and the tests were conducted from the least to most stressful (for the execution order of behavioral tests, see suppl. Fig. S2). Prior to testing, all animals were assessed using the SHIRPA protocol (Rogers et al., 1997). Mice were acclimated in the behavioral room for 30 min before the examination. Testing chambers were cleaned with 2% Derquim solution between animals to avoid olfactory cues. All behavioral tasks and quantification analyses were performed by researchers who were blinded to the mice’s treatment groups. Behavioral tests were carried out to evaluate phenotypes associated with positive (open field, self-grooming, startle reflex and prepulse inhibition, rotarod and hotplate tests), negative (three-chamber and tube test) and cognitive (novel object recognition and y-maze tests) symptoms (for the precise protocols, see suppl. mat. 1.3). Mice were sacrificed on PD75 and their brains were extracted.

### 2.4 Biochemical analysis

The PFC was dissected using a stainless brain matrix (Agnthos, 1mm) and snap-frozen in liquid nitrogen following cervical dislocation on PD75. Total protein was extracted from PFC samples of the four experimental groups and both sexes (n = 3-5 mice per group and sex). Briefly, PFC samples were lysed in 150-200 µl of lysis buffer, disaggregated using a syringe and then centrifuged at 10,000-17,000 x g at 4 °C for 10 minutes. Protein concentration was quantified using a BCA protein assay kit (23227, Thermofisher Scientific).

#### 2.4.1 Proteomic analysis

Proteomic analysis was carried out in the Proteomic services at IBiS. Proteins were reduced and cysteine residues blocked with 50 mM tris-(2-carboxyetyl) phosphine (TCEP, AB Sciex)) and cloroacetamide (CAA, Sigma-Aldrich, St. Louis, MO, USA)) for 30 minutes at 37°C with shaking. Proteolysis was carried out at 37 °C with trypsin (Promega, Fitchburg, WI, USA) at 1:20 (enzyme to substrate) at 37 °C overnight with agitation. The resulting peptide samples were then quenched with formic acid (pH 3) and desalted using a Stage-tip (Omix C18 tips, Agilent). Analysis of digested peptides was performed by LC-MS/MS with an Easy-nLC 1000 HPLC system (Thermo Fisher) coupled to a Q Exactive Plus Orbitrap mass spectrometer (Thermo Fisher). LC separations were performed on C18 HPLC precolumn (75 μm x 2 cm) and Easy Spray HPLC column (75 μm x 25 cm, 2 μm, 100 Å). Gradient elution was performed with a binary system consisting of (A) 0.1% aqueous formic acid and (B) 0.1% formic acid in CH3CN during 120 minutes from 5 – 40 % gradient of B with a flow rate of 250 nl/minute. A MS survey scan was obtained using a top 15 method with 100–2000 m/z mass range. The MS/MS Spectra Acquisition was obtained using Higher-energy Collisional Dissociation (HCD). An isolation mass window of 2 m/z was used for the precursor ion selection, and normalized collision energy of 27 % was used for fragmentation. Ten second duration was used for the dynamic exclusion.

Protein was quantified by spectrometry-based label-free identification. Mass spectrometry data analysis was performed through Proteome Discoverer (Thermo Fisher Scientific) with search an Uniprot database for Mus Musculus with a false discovery rate FDR less than 1%. Ratios for each experimental group were calculated relative to the control group. The results were analyzed using Ingenuity Pathway Analysis (IPA) software (Qiagen).

#### 2.4.2 Western blot

Equal amounts of protein (25-40 µg) were denatured by boiling in Laemmli buffer for 5- 10 minutes and then separated by SDS-PAGE using 6-12% polyacrylamide gels. Proteins were transferred to a PVDF membrane (Millipore). The membrane was blocked with 5% non-fat milk or BSA in Tris-buffered saline with Tween 20 (TBS-T) and incubated with the following primary antibodies: mouse anti-Reelin (1:1000, MAB5364, Sigma-Aldrich), rabbit anti-Dab1 (1:500, #3328, Cell Signaling) or mouse anti-β-Actin (1:1000, A5316, Sigma-Aldrich). Immunoreactivity was detected using the appropriate secondary antibodies conjugated with horseradish peroxidase (HRP) (1:5000): goat anti-mouse-HRP (NA931, Amersham) or donkey anti-rabbit-HRP (NA934, Amersham). Detection was performed using Clarity ECL Substrate (Bio-Rad) on a Chemidoc Touch Imaging System (Bio-Rad).

### 2.5 Histology

#### 2.5.1 Immunofluorescence

Animals were anesthetized with a lethal dose of thiobarbital (Braun) and perfused with PBS (n = 3-4 per group and sex). Brains were extracted, fixed overnight at 4 °C in 4% PFA and then immersed in 30% sucrose solution in PBS for 48 hours. Coronal sections of 25 µm thickness were obtained using a cryostat (Leica). For antigen retrieval, sections were incubated in a sodium citrate solution (10 mM, pH 6), and non-specific binding was blocked with 10% goat serum in PBS-T for 1 h. Sections were then incubated overnight at 4 °C with primary antibody mouse anti-Reelin (1:500, MAB5364, Sigma-Aldrich) and secondary antibody goat anti-mouse Alexa 594 (1:500, A-21203, Life Technologies). Sections were counterstained with DAPI and mounted on Superfrost-Plus slides with DAKO mounting media (Agilent). Images were acquired in a Thunder microscope (Leica) with a 20x objective. All images were analyzed using the ImageJ software (National Institute of Health). Density of Reelin+ cells was automatically quantified using a minimum of 4 slices per mice. The quantification of positive cells in the prefrontal cortex was performed by averaging three randomly selected fields within the prelimbic area. Cortical thickness measurement was carried manually out using the DAPI staining in the same mice in the primary somatosensory area.

#### 2.5.2 Golgi-Cox staining

Golgi-Cox staining was performed as described previously (Glaser & Van Der Loos, 1981). The Golgi-Cox solution (1% K_2_Cr_2_O_7_, 1% HgCl_2_, 0.8% K_2_CrO_4_) was prepared 5 days prior to use. Animals were anesthetized, decapitated, and the brains were extracted and hemisected (n = 5-16 per group). The hemispheres were immersed in Golgi-Cox solution for 48 h, after which the solution was refreshed, and the brains were incubated for 3 weeks. Coronal sections of 200 µm were cut using a vibratome (Leica) and incubated with 14-16% ammonium hydroxide for 1 hour. Sections were then treated with sodium thiosulfate for 7 minutes, dehydrated through an ethanol series (50-100%) and cleared with xylene. Sections were mounted in Superfrost-Plus slides with DPX mounting media. Z-stacks images of a 0.75 μm step size were acquired in a BX-61 bright- field microscope (Olympus) with a 100x objective. Secondary dendritic segments with a minimum length of 10 µm were selected, and dendritic spines were manually counted using ImageJ software (National Institute of Health). Spine density was calculated for each segment and averaged for each neuron.

### 2.6 RT-qPCR

Total RNA was extracted from homogenized maternal spleen or embryo (with placenta) samples with the RNeasy mini kit (74104, Qiagen). For each sample (n = 2 dams and n = 6 embryos per group), 1 µg of total RNA was used for reverse transcription to cDNA (qScript cDNA supermix, 95048-100, Quantabio), according to the manufacturer protocols. qPCR was performed with iTaq Universal Probes Supermix (Bio-Rad) and Taqman probes (IL-6: Mm00446190_m1; TNFα: Mm900443258_m1; GADPH: Mm99999915_g1, Thermofisher Scientific) on a ViiA 7 Real-Time PCR system (Thermofisher Scientific). Each sample was analyzed in duplicate.

### 2.7 Statistical analysis

Data are presented as the mean ± standard error of the mean (SEM). Statistical procedures were performed using GraphPad Prism™ 8 software (Graph Pad Software Inc., San Diego, CA, USA). Significance was defined as p < 0.05. Appropriate statistical tests, including three-way ANOVA, two-way ANOVA, and unpaired Student’s t-test, were performed. Post-hoc multiple comparisons were carried out using Tukey’s or Sidak’s correction, as applicable.

## 3. Results

### 3.1 Behavioral characterization of the dual-hit mouse model

The diagnosis of SCZ relies on assessing positive, negative, and cognitive symptoms. To investigate whether prenatal Poly(I:C) infection and postnatal THC exposure cause SCZ-related deficits, a behavioral battery was conducted in the model. Tests were performed to evaluate the face validity of the model.

#### 3.1.1 Male mice in the dual-hit model exhibit enhanced stereotyped behavior

The effects of prenatal Poly(I:C) and postnatal THC, either alone or in combination, on locomotion and anxiety were evaluated using the open field test. While THC does not affect these parameters after an extended drug-free period (Harte & Dow-Edwards, 2010; Kasten et al., 2019), acute THC exposure can alter anxiety and locomotion in mice (Kasten et al., 2019).

In the open field test, a three-way ANOVA revealed significant main effects of sex and THC on both the total distance traveled (Sex: F(1, 99) = 6.069, p < 0.05; THC: F(1, 99) = 14.21, p < 0.001) and the mean velocity (Sex: F(1, 99) = 6.142, p < 0.05; THC: F(1, 99) = 14.51, p < 0.001) (Fig. 2A). In male mice, a two-way ANOVA revealed a significant main effect of THC treatment on both the total distance traveled (F(1, 52) = 10.27, p < 0.01) and the mean velocity (F(1, 52) = 10.39, p < 0.01) in the open field test. Post-hoc comparisons showed that the Saline + THC group exhibited reduced locomotor activity and mean velocity compared to the Poly(I:C) + Vehicle group (p < 0.05). Similarly, the Poly(I:C) + THC group also displayed a significant decrease in mean velocity relative to the Poly(I:C) +Vehicle group (p < 0.05). Additionally, there was a trend toward decreased distance traveled in the Poly(I:C) + THC group compared to the Poly(I:C) + Vehicle group (p = 0.051). In female mice, a two-way ANOVA revealed a significant main effect of THC treatment on both total distance traveled (F(1, 47) = 5.363, p < 0.05) and mean velocity (F(1, 47) = 5.516, p < 0.05) in the open field test. Although post-hoc analyses did not detect statistically significant pairwise differences between groups, a general reduction in locomotor activity was observed in THC-treated groups, suggesting a trend toward THC-induced hypoactivity in females. Overall, these findings suggest that THC impairs locomotion regardless of prior immune activation, although the combination with Poly(I:C) may exacerbate the effect.

**Figure 2.**
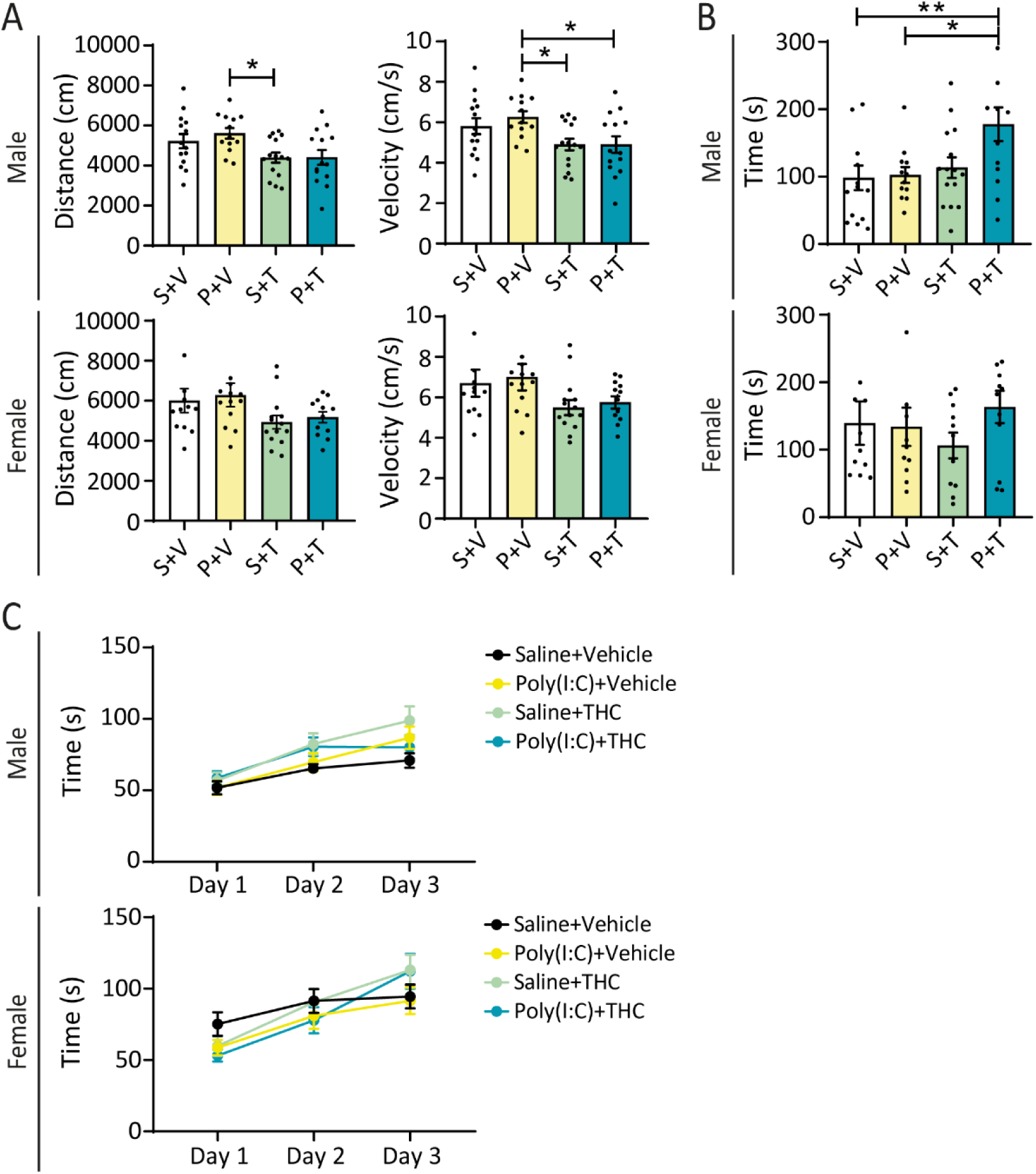
Poly(I:C) + THC male mice display increased stereotyped behavior. **A,** Open field test. Total distance traveled and velocity (n = 12-15 per treatment and sex) (two-way ANOVA with Tukey’s post-hoc analysis; *p < 0.05). **B,** Self-grooming time (n = 12-15 per treatment and sex) (two-way ANOVA with Tukey’s post-hoc analysis; *p < 0.05; **p < 0.01). **C,** Rotarod test. Latency to fall (n = 12-15 per treatment and sex) (repeated-measures three-way ANOVA with Tukey’s post-hoc analysis). Abbreviations: Saline + Vehicle (S+V), Poly(I:C) + Vehicle (P+V), Saline + THC (S+T), Poly(I:C) + THC (P+T).

Also regarding positive symptoms, the self-grooming test was used to study stereotyped behavior (Morrens et al., 2006). Several MIA models have reported an increased grooming time (Schwartzer et al., 2013; Haddad et al., 2020; Lan et al., 2023). We evaluated the possible enhanced effect of the dual-hit paradigm in stereotypy in the model. In the self-grooming test, a three-way ANOVA revealed a significant effect of Poly(I:C) (F(1, 90) = 4.183, p < 0.05) and a trend toward an interaction effect between Poly(I:C) and THC (p = 0.070) (Fig. 2B). In males, a two-way ANOVA showed significant effects of both THC (F(1, 49) = 7.196, p < 0.01) and Poly(I:C) (F(1, 49) = 4.361, p < 0.05).

Post-hoc analyses indicated increased grooming time in the Poly(I:C) + THC group compared to the control group (p < 0.01) and the Poly(I:C) + Vehicle group (p < 0.05). Additionally, there was a trend toward increased grooming time in the Poly(I:C) + THC group compared to the Saline + THC group (p = 0.063). In females, no significant differences were observed in the two-way ANOVA. These results indicate that the combination of Poly(I:C) and THC exerts a synergistic effect on self-grooming time in males.

On the other hand, cannabis exposure affects motor learning and performance (Prashad & Filbey, 2017) as well as movement speed and balance (Hitchcock et al., 2021). In the rotarod test, a repeated-measures three-way ANOVA revealed a significant main effect of day (F(1.63, 84.93) = 41.32, p < 0.0001) and a significant day × Poly(I:C) x THC interaction (F(2, 104) = 4.193, p < 0.05). Post hoc comparisons showed a general trend of improved motor performance in all treatment groups (Poly(I:C) alone, THC alone, and the combined treatment), reflected by increased latency to fall compared to control animals. However, no statistically significant differences were detected between treatment groups within individual testing days (Fig. 2C). In female mice, the analysis revelaed a significant main effect of day (F(1.81, 85.12) = 47.44, p < 0.0001) and a significant day x THC interaction (F(2, 94) = 6.736, p < 0.01). However, post-hoc comparisons did not detect statistically significant differences between treatment groups at any individual time point.

There were no significant differences between groups or sexes in heat sensitivity (hotplate) (suppl. Fig. S3). In the startle reflex and prepulse inhibition test, a main effect of stimulus intensity was observed in both the prepulse inhibition phase (F(1.47, 47.08) = 33.02, p < 0.0001) and the acoustic startle response phase (F(2.33, 74.53) = 169.2, p < 0.0001). However, no main effects of Poly(I:C), THC or their interaction were detected (suppl. Fig. S4).

#### 3.1.4 The dual-hit model impairs social novelty in both sexes and reduces dominance in male mice

SCZ patients display negative symptoms such as social withdrawal and anhedonia (Chesworth & Karl, 2017) which are modeled in animals through social tests like the three-chamber test or the tube test. In the social preference phase of the three-chamber test, no significant effects were detected in the three-way ANOVA in the social index (Fig. 3A). However, in the social novelty phase, the analysis showed significant main effects of both Poly(I:C) (F(1, 99) = 7.125, p < 0.01) and THC (F(1, 99) = 11.39, p < 0.01) in the memory index (Fig. 3B). As sex showed no significant main effect nor any significant interactions with the other factors, data were pooled across sexes. A two-way ANOVA revealed significant main effects of both prenatal Poly(I:C) (F(1, 103) = 7.482, p < 0.01) and postnatal THC (F(1, 103) = 11.86, p < 0.001) on the memory index. Post-hoc analysis showed a significant reduction in the memory index in the Poly(I:C) + THC group compared to controls (p < 0.001), as well as in the Saline + THC group (p < 0.01) and the Poly(I:C) + Vehicle group (p < 0.05), indicating that both treatments independently and additively impaired recognition memory performance.

**Figure 3.**
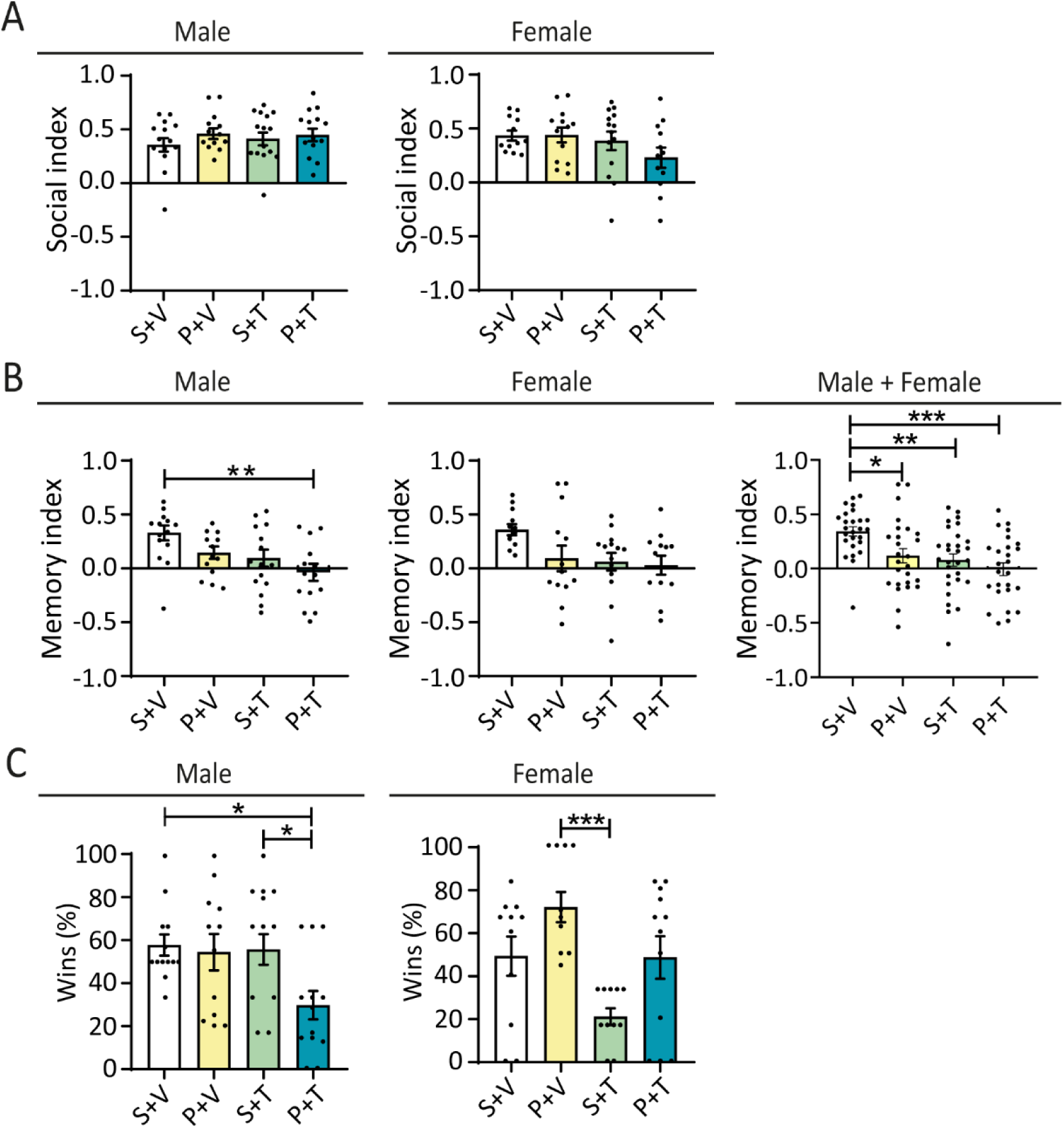
The dual-hit paradigm impairs social novelty in both sexes and dominance in male mice. **A,** Three-chamber test. Social index in the social preference phase (n = 12-15 per treatment and sex) (two-way ANOVA with Tukey’s post-hoc analysis). **B,** Three-chamber test. Memory index in the social novelty phase (n = 12-15 per treatment and sex) (two-way ANOVA with Tukey’s post-hoc analysis; **p < 0.01). **C,** Tube test. Percentage of wins (n = 12-15 per treatment and sex) (two-way ANOVA with Tukey’s post-hoc analysis; *p < 0.05, ***p < 0.001). Abbreviations: Saline + Vehicle (S+V), Poly(I:C) + Vehicle (P+V), Saline + THC (S+T), Poly(I:C) + THC (P+T).

Regarding social dominance, a three way ANOVA revealed a significant main effect of THC on the percentage of wins in the tube test (F(1, 90) = 13.98, p < 0.001), as well as a significant sex x Poly(I:C) interaction (F(1, 90) = 14.42, p < 0.001) (Fig. 3C). Analysis of paired treatments is available in suppl. Fig. S5. In male mice, a two-way ANOVA showed a significant main effect of Poly(I:C) on the percentage of wins (F(1, 49) = 4.483, p < 0.05), along with a trend toward a main effect of THC (p = 0.058). Post-hoc comparisons showed that Poly(I:C) + THC male mice displayed a significantly lower proportion of wins compared to controls (p < 0.05) and Saline + THC animals (p < 0.05). Additionally, a trend toward reduced dominance was observed when comparing Poly(I:C) + THC males to Poly(I:C) + Vehicle animals (p = 0.077), suggesting a potential synergistic effect of the two exposures on social dominance behavior. Regarding females, a two-way ANOVA revealed significant main effects of both Poly(I:C) (F(1, 41) = 10.01, p < 0.01) and THC (F(1, 41) = 10.49, p < 0.01). Post-hoc comparisons showed that Saline + THC females exhibited a significantly lower proportion of wins compared to the Poly(I:C) + Vehicle group (p < 0.001). Additionally, there was a trend toward reduced dominance in the Saline + THC group compared to both the controls (p = 0.079) and the Poly(I:C) + THC group (p = 0.072), suggesting that THC alone may impair social dominance behavior in females, independently of MIA.

#### 3.1.6 Poly(I:C) treatment alone impairs recognition memory in female mice

THC exposure in rodent models causes short-term recognition memory alterations (Kirschmann et al., 2017; Abush et al., 2018), as well as prenatal Poly(I:C) infection (Ratnayake et al., 2012; Gray et al., 2019; Sánchez-Hidalgo et al., 2022). In the novel object recognition test, regarding the memory index, a three-way ANOVA revealed a significant main effect of Poly(I:C) (F(1, 78) = 8.708, p < 0.01), as well as a significant sex x THC interaction (F(1, 78) = 5.139, p < 0.05). Additionally, a trend toward a main effect of THC was observed (p = 0.072). In males, a two-way ANOVA showed no significant differences between groups. However, in females, there was a main effect of Poly(I:C) (F(1, 36) = 6.467, p < 0.05) and THC (F(1, 36) = 8.294, p < 0.01). Post-hoc analysis revealed a reduced memory index in Poly(I:C) + Vehicle mice compared to controls (p < 0.05), Saline + THC (p < 0.01) and Poly(I:C) + THC (p < 0.05) mice (Fig. 4). There were no differences in spontaneous alternations in the y-maze test (suppl. Fig. S6). These findings suggest that prenatal immune activation with Poly(I:C) selectively impairs object recognition memory in female mice, while no significant effects were observed in males.

**Figure 4.**
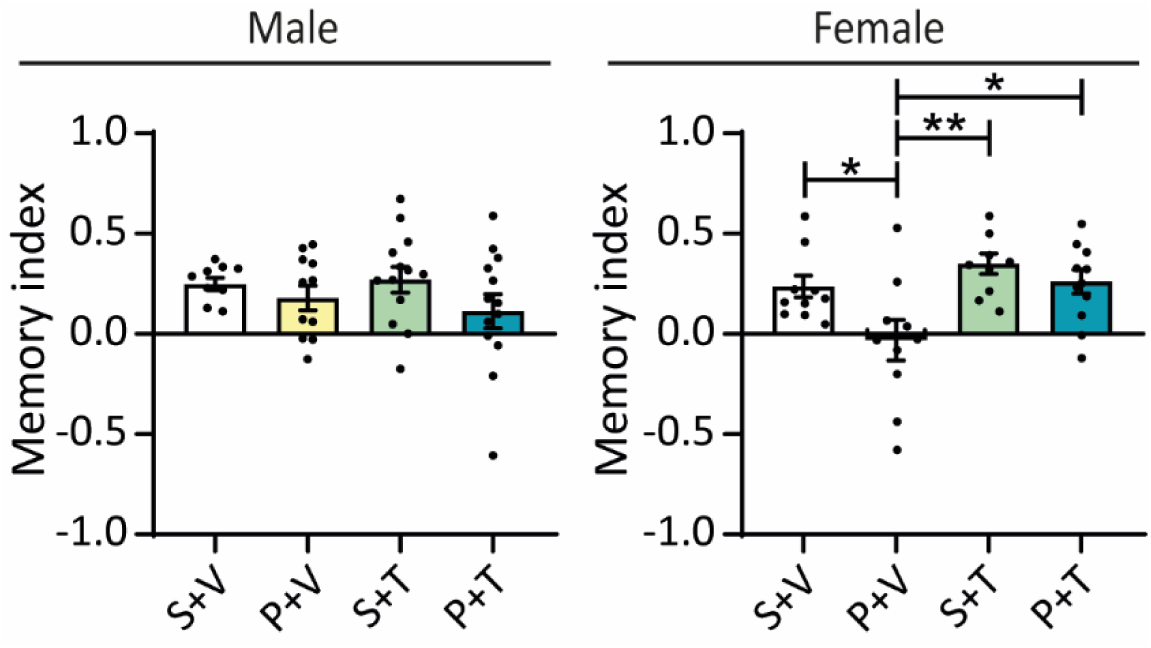
Recognition memory is altered by Poly(I:C) treatment alone in female mice. Novel object recognition test. Memory index (n = 12-15 per treatment and sex) (two-way ANOVA with Tukey’s post-hoc analysis; *p < 0.05, **p < 0.01). Abbreviations: Saline + Vehicle (S+V), Poly(I:C) + Vehicle (P+V), Saline + THC (S+T), Poly(I:C) + THC (P+T).

### 3.2 The Reelin signaling pathway is altered in the dual-hit mouse model

#### 3.2.1 Proteomic analysis

PFC samples from five mice per treatment group (Poly(I:C) and/or THC) and four control mice (Saline + Vehicle), including both sexes, were analyzed using label-free quantitative proteomics (Fig. 5). The raw data from this analysis is available as Supplemental Data (suppl. mat. 3). This analysis served to identify proteins significantly altered (p < 0.05) in dual-hit male and female mice (Fig. 5A, B). A total of 375 altered proteins were detected in male mice and 447 in female mice compared to controls (Fig. 5A).

**Figure 5.**
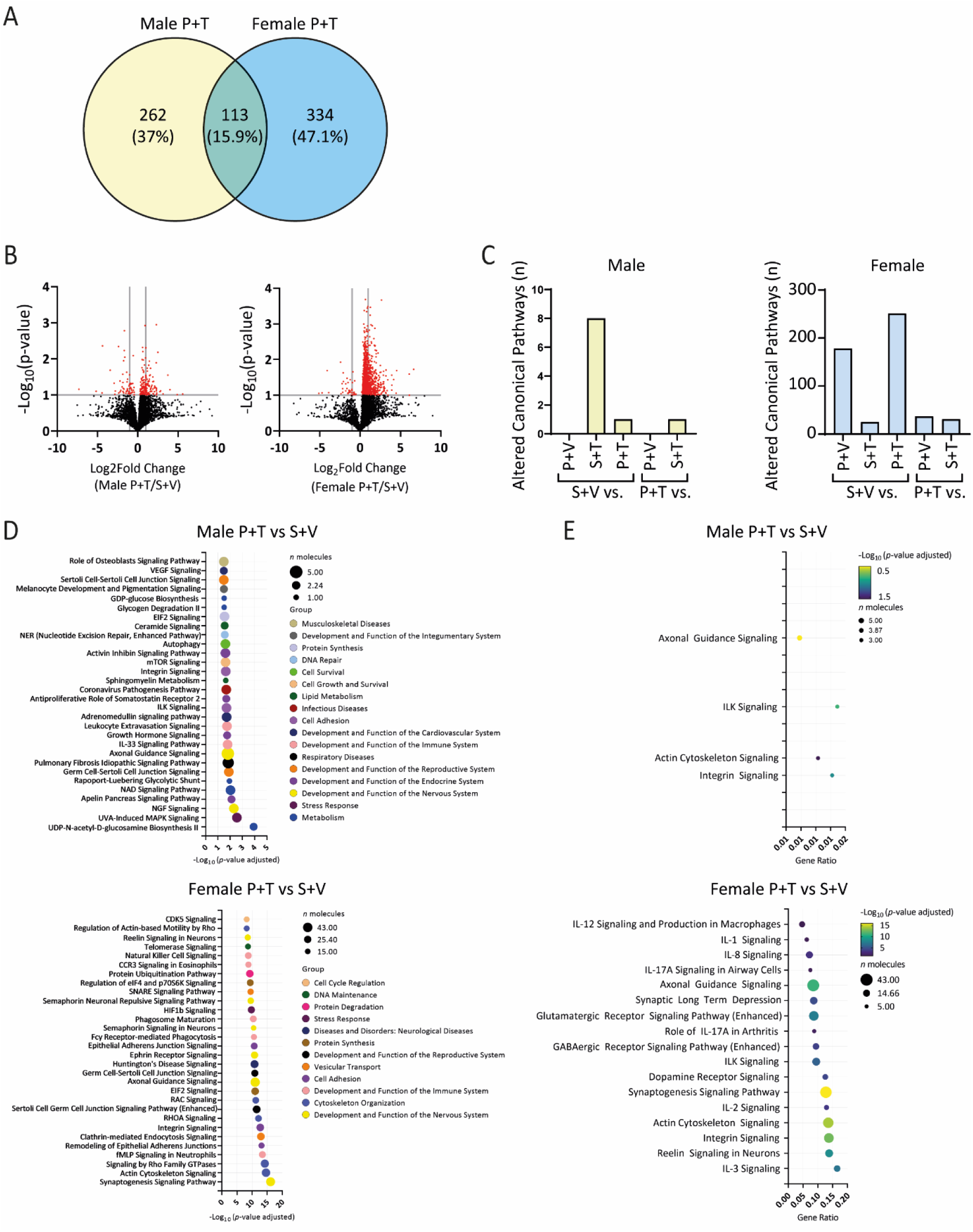
Quantitative proteomics analysis of PFC samples from Poly(I:C)- and/or THC-treated mice. **A,** Venn diagram showing the number and percentage of proteins altered after Poly(I:C) + THC treatment in males (yellow) and females (blue) compared to controls, including those commonly affected in both sexes. **B,** Volcano plot illustrating the distribution of differentially expressed proteins based on log2(Fold Change) and -log10(p-value) for the Poly(I:C) + THC male vs. Poly(I:C) + THC female comparison, normalized to their respective controls. Proteins with significant differential expression (p < 0.05) are highlighted in red. **C,** Count of significantly altered canonical pathways (p < 0.05) identified in each indicated comparison for males and females. **D,** Ingenuity Canonical Pathways enrichment analysis depicting significantly differentially expressed pathways in Poly(I:C) + THC-treated males (upper panel) and females (bottom panel), normalized to their respective controls. Dot size corresponds to the number of altered proteins within each pathway (n molecules). **E,** Dot plot quantifying the number of significantly altered proteins per pathway set and the total dataset (Gene Ratio) in males (upper panel) and females (bottom panel), normalized to their respective controls. Color intensity represents the adjusted p-value (-log10), while dot size indicates the number of altered proteins within each pathway. Abbreviations: Saline + Vehicle (S+V), Poly(I:C) + Vehicle (P+V), Saline + THC (S+T), Poly(I:C) + THC (P+T).

Notably, 113 proteins were common to both conditions, suggesting a core set of proteins consistently expressed under the dual-hit paradigm. Furthermore, 262 proteins were uniquely altered to male mice, while 334 were exclusive to females (the list of altered proteins in males, females, and common to both is available in suppl. mat. 4). These findings indicate a sex-specific protein signature of the dual-hit paradigm and highlight potential therapeutic targets for further investigation. A higher proportion of altered proteins was observed in female dual-hit mice compared to their controls (Fig. 5B). These results suggest that the dual-hit paradigm induces a more extensive protein expression response in females than in males, highlighting potential sex-specific mechanisms underlying this condition. A similar pattern was also observed in the analysis of altered canonical pathways, showing a higher number in female dual-hit mice compared to their controls and Saline + THC male mice (Fig. 5C). The 30 most significantly altered canonical pathways between the dual-hit groups and their respective controls are shown in Fig. 5D. Among the pathways most relevant to SCZ, significant alterations were observed in interleukin (IL) signaling -specifically IL-1, IL-2, IL-3, IL-8, IL-12 and IL-17A- as well as in synaptogenesis, axonal guidance, actin cytoskeleton and integrin signaling, long term depression, and glutamatergic, GABAergic, and dopamine receptor pathways (Fig. 5E). Notably, the Reelin signaling pathway in neurons emerged as particularly noteworthy due to its relevance in the context of SCZ. This pathway was significantly altered in Saline + THC males compared to their controls (suppl. Fig. S7), and across all treatment groups (Poly(I:C) + Vehicle, Saline + THC y Poly(I:C) + THC) when compared to their respective female controls, highlighting its potential role within the dual-hit paradigm. Similar results were observed in the analysis comparing Saline + THC and control mice (suppl. Fig. S6). Briefly, a total of 293 proteins were found to be altered uniquely in male Saline + THC mice, and 303 in female mice, with 91 altered proteins common to both groups. The most significantly altered canonical pathways were associated with GABAergic and glutamatergic signaling, Reelin signaling in neurons, as well as ILK, actin cytoskeleton, IL-2, and integrin signaling.

#### 3.2.2 Reelin and Dab1 expression levels

The alteration in the Reelin signaling pathway was confirmed via Western blot analysis in PFC samples by evaluating the expression levels of Reelin, its N-R2 fragment and Dab1 (Fig. 6A). Regarding Reelin expression, a three-way ANOVA reveled a significant main effect of sex (F(1, 16) = 5.408, p < 0.05), as well as a significant interaction between Poly(I:C) and THC (F(1, 16) = 10.21, p < 0.01). Additionally, there were trends toward significance for the main effects of THC (p = 0.051) and Poly(I:C) (p = 0.072), as well as for the sex x Poly(I:C) interaction (p = 0.094). In light of the significant sex effect, further analyses were conducted separately for male and female subjects using two-way ANOVAs.

**Figure 6.**
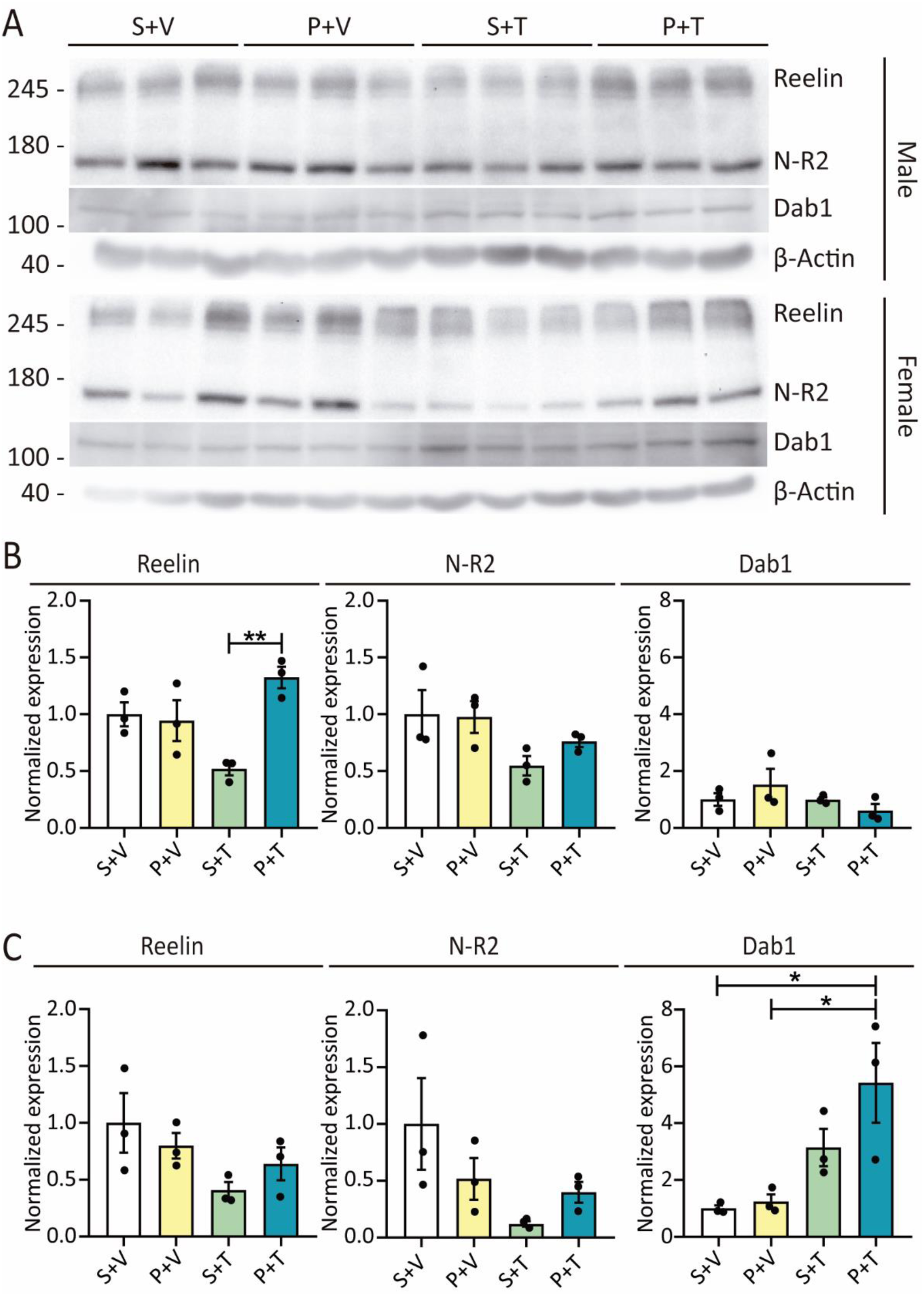
Reelin and Dab1 expression are altered in the model. **A,** Western blot experiments from PFC lysates of S+V, P+V, S+T and P+T male and female mice. **B,** Normalized expression of Reelin, N-R2 fragment and Dab1 (male mice) (two-way ANOVA with Tukey’s post-hoc analysis; **p < 0.01). **C,** Normalized expression of Reelin, N-R2 fragment and Dab1 (female mice) (two-way ANOVA with Tukey’s post-hoc analysis) (n = 3 per treatment and sex). Abbreviations: Saline + Vehicle (S+V), Poly(I:C) + Vehicle (P+V), Saline + THC (S+T), Poly(I:C) + THC (P+T).

In males, the analysis showed a main effect of Poly(I:C) (F(1, 8) = 10.07, p < 0.05) as well as a significant interaction between Poly(I:C) and THC (F(1, 8) = 13.27, p < 0.01) (Fig. 6B). Post-hoc analysis reported that Reelin expression was decreased in Saline + THC male mice compared to the dual-hit group (p < 0.01). In females, a two-way ANOVA revealed a trend toward a main effect of postnatal THC treatment (p = 0.051), with no significant interaction between prenatal and postnatal treatments (Fig. 6C). Although post-hoc comparisons did not reach statistical significance, THC-exposed groups showed a general reduction in Reelin expression compared to controls, suggesting a consistent downward trend associated with postnatal THC exposure.

Regarding N-R2 fragment expression, a three-way ANOVA reported significant main effects of sex (F(1, 16) = 5.542, p < 0.05) and THC (F(1, 16) = 9.872, p < 0.01). Additionally, a trend toward a Poly(I:C) x THC interaction was observed (p = 0.077), suggesting a possible combined influence of prenatal and postnatal hits. In males, there was a main effect of THC (F(1, 8) = 5.934, p < 0.05), indicating an overall influence of THC in N-R2 expression (Fig. 6B). However, post-hoc comparisons did not identify statistically significant pairwise differences between specific groups. Despite this, a general reduction in N-R2 expression level was observed across both THC-treated groups compared to controls. In females, a two-way ANOVA revealed a trend toward a main effect of THC (p = 0.059) (Fig. 6C). Although post-hoc comparisons did not reach statistical significance, a consistent pattern of reduced N-R2 expression was observed across all treated groups compared to controls. Therefore, a reduction in both full-length Reelin and its N-R2 fragment levels was observed, rather than an accumulation of the fragment. This suggests that the decrease in Reelin expression is unlikely to be attributed to enhanced proteolytic processing of its N-terminal region.

With respect to Dab1 expression, a three-way ANOVA revealed significant main effects of sex (F(1, 16) = 15.51, p < 0.01) and THC (F(1, 16) = 10.05, p < 0.01), as well as a significant sex x THC interaction (F(1, 16) = 18.20, p < 0.001). These findings suggest that the effect of THC differs between sexes, supporting separate analyses by sex. In males, a two-way ANOVA did not show any effects (Fig. 6B). However, in females, there was a main effect of THC (F(1, 8) = 16.17, p < 0.01), and post-hoc analysis revealed an increased Dab1 expression in dual-hit mice compared to controls (p < 0.05) and Poly(I:C) + Vehicle mice (p = 0.05) (Fig. 6C). Dab1 expression has been utilized as an indicator of Reelin signaling activity, given that Reelin binding to its receptors, ApoER2 and VLDLR, triggers Dab1 phosphorylation and the initiation of downstream signaling cascades (Ogino et al., 2017). Therefore, the observed increase in Dab1 levels is consistent with the reduction in Reelin expression.

In summary, the non-significant reduction in expression levels of both full-length Reelin and its N-R2 fragment in the Saline + THC group in males, and across all three experimental groups in females, and the significantly altered Dab1 expression levels in females, collectively support the disruption of the Reelin signaling pathway observed in the proteomic study.

#### 3.2.3 Reelin+ cell density is reduced in the dual-hit female mice

The density of Reelin+ cells in the prelimbic area of the PFC was quantified via immunofluorescence. A three-way ANOVA revealed significant main effects of sex (F(1, 23) = 14.49, p < 0.001) and THC (F(1, 23) = 19.71, p < 0.001), as well as a significant sex x Poly(I:C) interaction (F(1, 23) = 7.017, p < 0.05). Additionally, a trend toward a three-way interaction (sex x Poly(I:C) x THC) was observed (F(1, 23) = 3.374, p = 0.079). Based on these findings, separated two-way ANOVAs were conducted for each sex to further examine the effects of prenatal and postnatal treatments.

In male mice, there was a main effect of THC (F(1, 12) = 5.534, p < 0.05), and post-hoc analysis showed a non-significant decrease in the Saline + THC mice compared to controls (p = 0.087). In contrast, in female mice, there was a main effect of both Poly(I:C) (F(1, 11) = 9.899, p < 0.01) and THC (F(1, 11) = 16.30, p < 0.01), and post-hoc analysis revealed a reduction in Poly(I:C) + THC compared to controls (p < 0.01) and Poly(I:C) treatment alone (p < 0.05). Moreover, the dual-hit group showed a marginally significant decrease in Reelin+ cell density compared to Saline + THC group (p = 0.053) (Fig. 7). These results are consistent with the findings from the Western blot experiments, as a reduced density of Reelin+ cells was observed in the same groups where the lower expression levels of full-length Reelin protein were detected.

**Figure 7.**
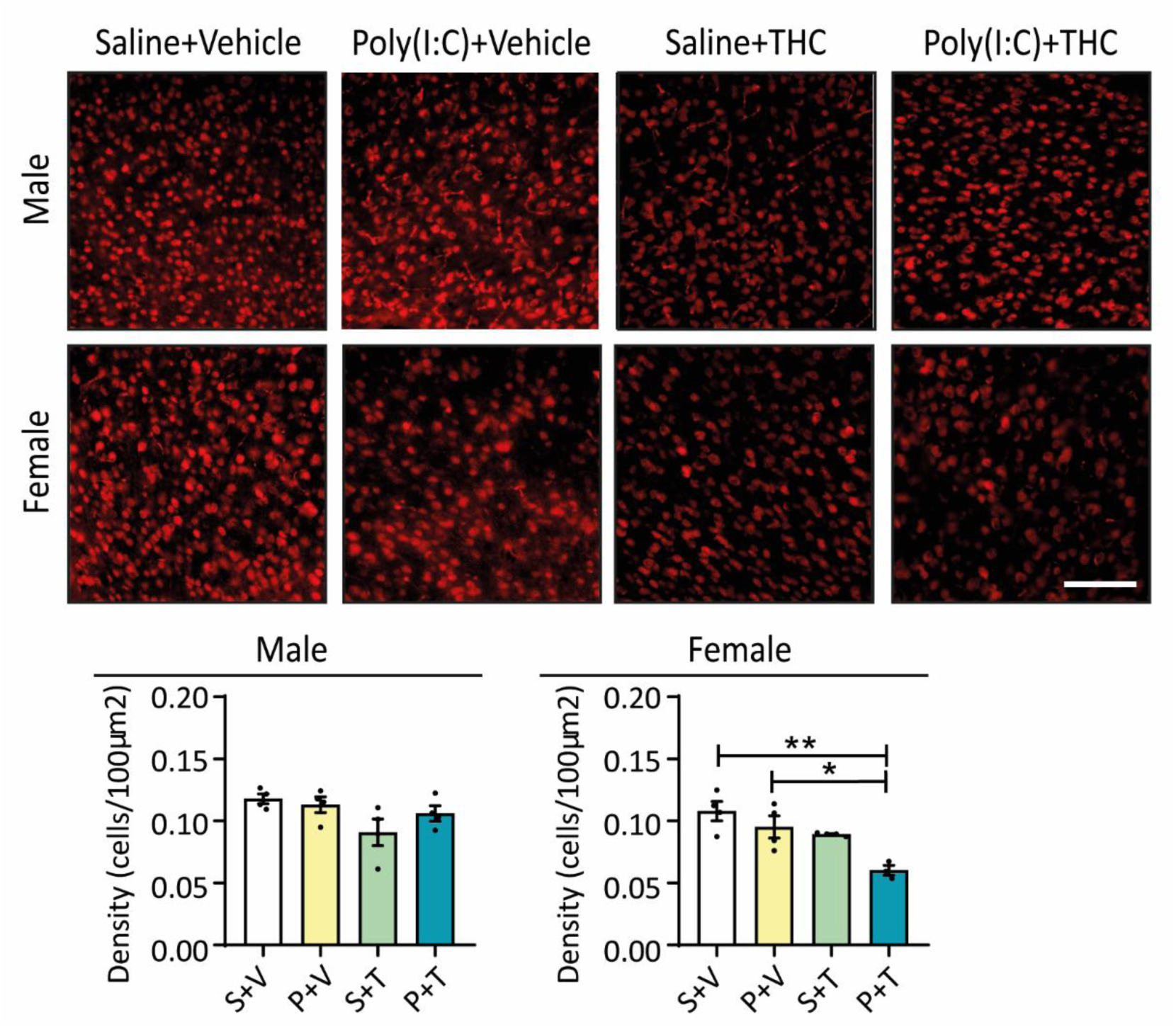
Reelin+ cell density is reduced in the dual-hit mice (two-way ANOVA with Tukey’s post-hoc analysis; *p < 0.05; **p < 0.01) (n = 3-4 per treatment and sex). Abbreviations: Saline + Vehicle (S+V), Poly(I:C) + Vehicle (P+V), Saline + THC (S+T), Poly(I:C) + THC (P+T). Scale: 180µm.

### 3.3 Total cortical thickness is reduced by THC exposure

SCZ patients exhibit reduced cortical thickness across several brain regions, such as the frontal cortex (Nenadic et al., 2015; Yan et al., 2019). Similarly, MIA models also show alterations in cortical thickness (Smith et al., 2012; Guma et al., 2019; Baines et al., 2020; Vlasova et al., 2021), though results can vary based on the timing of injection and dosage. Additionally, cannabis use can impact cortical thickness in SCZ patients (Rais et al., 2010; Manza et al., 2020; Mashhoon et al., 2015).

Cortical thickness was manually quantified in the somatosensory area using DAPI staining in coronal brain slices. To evaluate potential influences of sex, prenatal treatment, and postnatal THC exposure, a three-way ANOVA was initially performed. Since no significant effects or interactions involving sex were detected, data from male and female animals were combined. A two-way ANOVA was then used to assess the effects of prenatal and postnatal THC treatment on the pooled dataset (Fig. 8).

**Figure 8.**
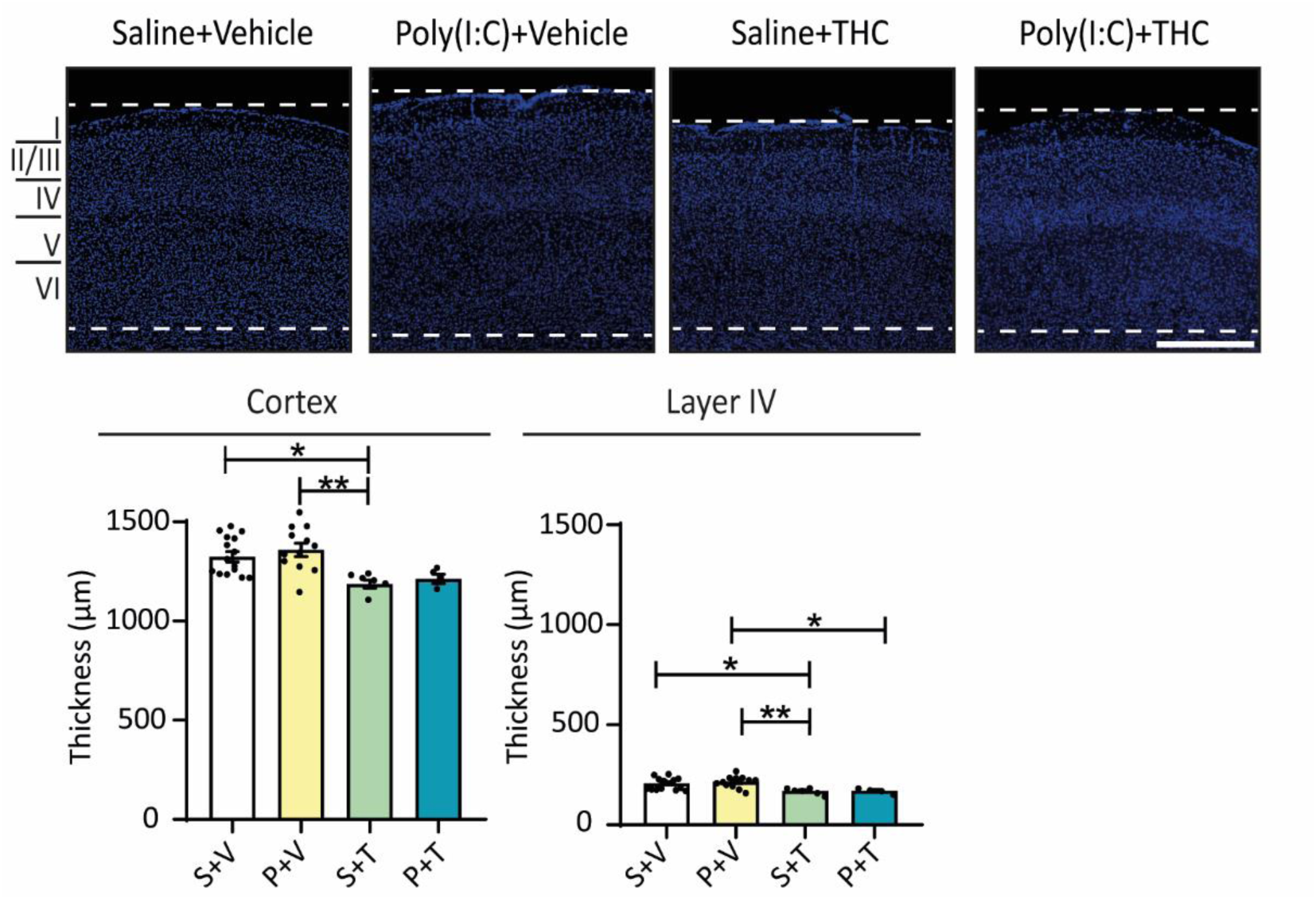
Total cortical thickness is reduced by THC treatment (total cortex and layer IV) (n = 4-15 per treatment) (two-way ANOVA with Tukey’s post-hoc analysis; *p < 0.05, **p<0.01). Abbreviations: Saline + Vehicle (S+V), Poly(I:C) + Vehicle (P+V), Saline + THC (S+T), Poly(I:C) + THC (P+T). Scale: 500µm.

A significant effect of THC in total cortical thickness was observed (F(1, 33) = 15.33, p < 0.001). Post-hoc analysis reported a significant reduction in Saline + THC mice compared to controls (p < 0.05) and Poly(I:C) + Vehicle (p < 0.01) groups. Poly(I:C) + THC showed a non-significant reduction in total cortex compared to Poly(I:C) treatment alone (p = 0.058). THC also showed a significant effect when cortical layers were analyzed separately, specifically in layer IV (F(1, 33) = 17.23, p < 0.001). Post-hoc analysis revealed a reduction in layer IV thickness in Saline + THC mice compared to controls (p < 0.05) and Poly(I:C) + Vehicle (p < 0.01). Dual-hit mice also showed a reduction compared to Poly(I:C) + Vehicle mice (p < 0.05). Taken together, the results suggest that postnatal THC exposure is the main factor contributing to the reduction in total cortical thickness and in layer IV, as significant effects were observed in both control and Poly(I:C)-exposed animals. Although the effect in the dual-hit group did not reach significance, a similar trend was noted.

### 3.4 Poly(I:C) + THC combination causes a decreased dendritic spine density in PFC and HP

Dendritic spine density was quantified in the prelimbic area of the PFC and CA1 of the HP using Golgi-Cox staining. A three-way ANOVA was conducted to assess the effects of sex, Poly(I:C) and THC treatment. As sex showed no significant main effect nor any significant interactions with the other factors, data were pooled across sexes. Subsequently, a two-way ANOVA (Poly(I:C) x THC treatment) was performed on the combined dataset.

In the CA1 region of the HP, the main effect of Poly(I:C) was observed on dendritic spine density (F(1, 36) = 9.514, p < 0.01). Post-hoc analysis indicated a significant reduction in dendritic spine density in the dual-hit group compared to controls (p < 0.05) (Fig. 9A). Similar findings were noted in the PFC, where a significant effect of Poly(I:C) was found (F(1, 36) = 16.47, p < 0.001). Post-hoc analysis revealed a significant decrease in dendritic spine density in the Poly(I:C) + Vehicle (p < 0.01) and dual-hit (p < 0.01) groups compared to controls and a non-significant decrease between Saline + THC and dual-hit group (p = 0.090) (Fig. 9B). Overall, these results indicate that dendritic spine density is affected by the initial impact, the Poly(I:C) prenatal infection, and is significantly further influenced by postnatal THC exposure. Thus, the dual-hit paradigm impacts dendritic spine density in both analyzed regions.

**Figure 9.**
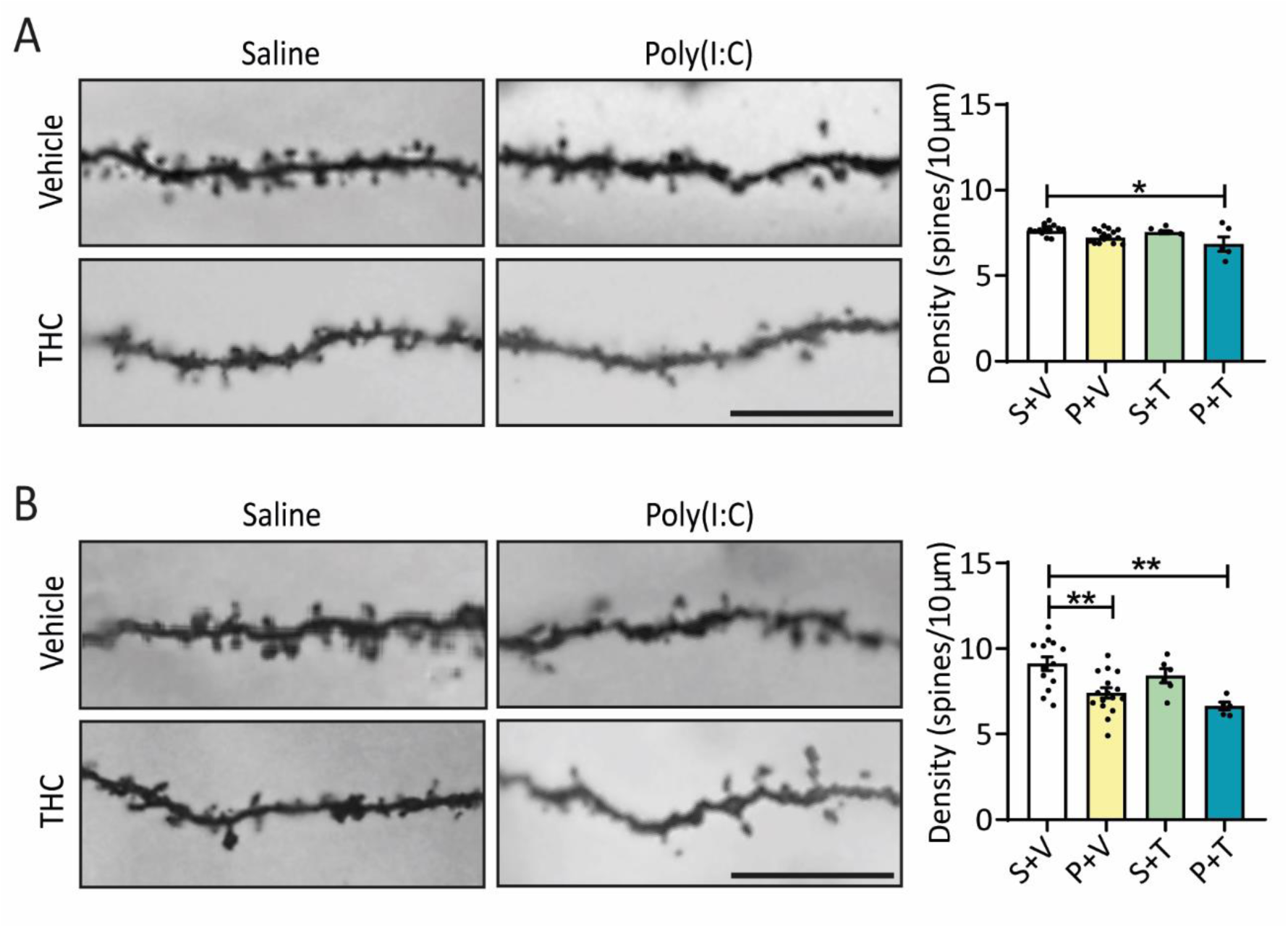
Poly(I:C) + THC treatment causes a decreased dendritic spine density in PFC and HP. **A,** Dendritic spine density in CA1 (HP) (n = 5-16 per treatment) (two-way ANOVA with Tukey’s post-hoc analysis; *p < 0.05). **B,** Dendritic spine density in PFC (n = 5-16 per treatment) (two-way ANOVA with Tukey’s post-hoc analysis; **p < 0.01). Abbreviations: Saline + Vehicle (S+V), Poly(I:C) + Vehicle (P+V), Saline + THC (S+T), Poly(I:C) + THC (P+T). Scale: 10µm.

## 4. Discussion

In this study, we established a dual-hit mouse model with MIA induced by Poly(I:C) on GD9 and postnatal THC exposure between PD55-PD60 to investigate the impact of THC as a secondary insult and assess disruptions in the Reelin signaling pathway. The model showed decreased Reelin expression, increased Dab1 expression, and reduced Reelin+ cell density, particularly in females, suggesting Reelin as a potential biomarker for SCZ and supporting the dual-hit hypothesis. The model also displayed behavioral phenotypes associated with positive symptoms in males and negative symptoms in both sexes. Additionally, THC exposure led to reduced cortical thickness and dendritic spine density in both PFC and HP. This model did not affect offspring viability and MIA was confirmed in all pregnant dams. Both male and female mice were included to assess potential sexual dimorphism (Ayesa-Arriola et al., 2020). We observed differences between sexes in body weight, behavior, proteomics, and Reelin signaling pathway alterations, emphasizing the importance of including both sexes in SCZ models.

Mouse models prenatally exposed to Poly(I:C) can exhibit alterations in stereotyped and social behavior, working memory, cognitive flexibility, and increased anxiety (Estes & McAllister, 2016; Gumusoglu et al., 2019). Our behavioral tests showed altered social behavior and stereotypy in dual-hit mice, consistent with previous studies indicating adolescence as a critical period for cannabis-induced symptoms (Chesworth & Karl, 2017).

Our dual-hit mouse model exhibited locomotor alterations, likely due to the acute THC effects, as the open field test was performed 24 hours after last injection (Kasten et al., 2019). In the tube test, Poly(I:C) + THC male mice showed reduced dominance. This effect was not detected in female mice, which may be due to the generally higher baseline dominance exhibited by males, making behavioral differences more detectable in this sex (Smith-Osborne et al., 2023; Karamihalev et al., 2020; Williamson et al., 2019; Van der Berg et al., 2015). No significant cognitive impairments were observed, possibly due to the timing of Poly(I:C) injection, which tends to cause more pronounced cognitive deficits when administered later in gestation (Chesworth & Karl, 2017; Gillespie et al., 2024). This is likely because MIA during advanced gestational stages has a more significant impact on the function and development of the GABAergic system, alterations of which have been linked to cognitive symptoms of the disorder (Gillespie et al., 2024).

Recent studies have highlighted Reelin as a potential key biomarker that contributes to the face validity of SCZ-related models, and most Reelin alterations are typically found in the PFC or HP of animal models (Sánchez-Hidalgo et al., 2022). Proteomics and IPA analysis in PFC samples revealed an altered Reelin signaling pathway in the majority of pairwise treatment comparisons, particularly in female mice. This was corroborated by Western blot analysis. It is noteworthy that there was an overexpression of Reelin in the dual-hit group rather than a reduction. In other dual-hit models, authors described a reversion of Poly(I:C)-induced phenotypes following adult THC exposure (Lecca et al., 2019). The prenatal impact could cause long-term alterations in the endocannabinoid system, which might subsequently be mitigated by the THC exposure. However, further analysis is required to test this hypothesis. Regarding female mice, a non-significant decrease in Reelin expression was observed across all three treatment groups, suggesting that THC exposure has a greater impact in Reelin expression compared to the prenatal Poly(I:C) injection. This decrease in Reelin expression could be due to increased proteolytic processing of its N-terminal end, leading to an accumulation of the N-R2 fragment (Jossin et al., 2004, 2007; Okugawa et al., 2020). However, the analysis of the N-R2 fragment revealed the same expression pattern as full-length Reelin, indicating that the reduction in full-length Reelin expression is not due to increased proteolytic processing of its N-terminal end.

On the other hand, the immunofluorescence study of Reelin+ cells showed a non-significant decrease in THC-treated male groups, as well as a significant reduction in all three experimental female groups compared to controls, with greater differences in the dual-hit group. This finding aligns with the Western blot results. Moreover, since the binding of Reelin to its receptors ApoER2 and VLDLR induces Dab1 phosphorylation and subsequent signaling cascades, some authors suggest that Dab1 concentration can be used as an indicator of Reelin signaling potency (Ogino et al., 2017). Dab1 levels were not altered in males, but a significant increase in its expression was observed in the dual-hit female group compared to their controls and the Poly(I:C) group. This increase in Dab1 levels is consistent with the reduction in Reelin, corroborating the disruption of the Reelin signaling pathway in the model.

The underlying neurobiological mechanisms involved in the interaction between prenatal infection and postnatal THC exposure are of significant interest for understanding the altered biological systems associated with the disease and remain incompletely characterized. Several authors have proposed various signaling pathways where the mechanisms of MIA and THC converge, focusing on communication between neurons, microglia, and astrocytes in response to this dual-hit combination (Murlanova & Pletnikov, 2023). Building on this hypothesis, we aim to highlight the potential role of the ECM as an additional convergent factor in neuron-glia communication, with a particular emphasis on the role of Reelin within the framework of the dual-hit hypothesis (Martín-Cuevas et al., 2023).

Reduced cortical thickness in various brain regions, such as the PFC, has been reported in individuals with SCZ through neuroimaging studies (Nenadic et al., 2015; Yan et al., 2019) and in patients with chronic cannabis use (Manza et al., 2020). In our model, a reduction in total cortical thickness and, particularly, in layer IV, was observed in the experimental groups treated with THC. Quantifying neuronal density in these layers would be necessary to determine whether the thinning of the affected layers results from neurodegeneration, neuronal death, or other alternative causes not implying death. On the other hand, a reduction in the density of dendritic spines in the pyramidal neurons of the PFC has been documented in individuals with SCZ (Garey, 2010). In our model, a significant reduction in dendritic spine density was observed in dual-hit mice. During prenatal stages, the primary source of Reelin is Cajal-Retzius cells located in the marginal zone of the cerebral cortex (Jossin, 2020; Hattori & Kohno, 2021). The observed impairment in dendritic spine density in the model may be attributed to reduced Reelin expression in Cajal-Retzius cells during prenatal stages as a consequence of MIA induced by Poly(I:C) injection on GD9, which is further exacerbated by postnatal THC exposure (Ogino et al., 2017).

This study presents several limitations that should be acknowledged. First, most of the experiments were conducted in PFC, which restricts the interpretation of the results to this specific brain region. Expanding the analyses to other relevant areas, such as the HP, would provide a more comprehensive understanding of the alterations observed in the model. Second, there are inherent limitations associated with the use of animal models of SZ.

Although mouse models allow for controlled and reproducible investigation of disease-related alterations, they cannot fully recapitulate the complexity of the disorder, particularly symptoms such as hallucinations, delusions, disorganized speech, and verbal learning deficits. Furthermore, behavioral tests were selected to assess potential phenotypes related to the positive, negative, and cognitive symptoms of SZ; however, it would be valuable to include additional paradigms targeting domains such as fear conditioning, spatial learning, or anxiety-related behaviors. Lastly, both the dosage and route of THC administration may significantly influence the observed outcomes. Investigating alternative administration routes, such as inhalation, could yield important insights into the translational relevance of the findings. These limitations should be taken into account when interpreting the results and drawing conclusions from the present study.

## 5. Conclusions

We established a dual-hit mouse model consisting of MIA with Poly(I:C) on GD9 and subsequent THC exposure during postnatal stages (PD55-60). Male mice subjected to the dual-hit paradigm exhibit stereotyped behavior and social impairments consistent with both positive and negative symptoms of SCZ and dual-hit female mice display social deficits. These findings support the face validity of the model. Our findings reveal that THC exposure leads to reduced Reelin expression and its N-R2 fragment in the PFC, particularly in females, as well as decreased Reelin+ cell density and increased Dab1 expression, confirming that the alteration of the Reelin signaling pathway is mainly caused by the adolescent THC exposure. THC also results in decreased cortical thickness. Additionally, dual-hit mice show reduced dendritic spine density in both the PFC and HP. We propose that this paradigm represents valid model for investigating the dual-hit hypothesis of SCZ and highlight Reelin as a potential key biomarker of the disease.

## Acknowledgements

This work was supported by the Consejería de Conocimiento y Universidades through project PID2019-109405RB-I00/AEI/10.13039/501100011033; the Andalusian Plan for Research, Development, and Innovation and ERDF/EU through project P20_00811; the Regional Ministry of Health (Junta de Andalucía) through project PI-0014-2022; the Instituto de Salud Carlos III (ISCIII) co-funded by the European Union, through projects AC23_2/00034 and PI22/01379, and unrestricted research funding from the Spanish Network for Research in Mental Health (CIBERSAM, G26). CM-C was supported by CIBERSAM (G26) and the Instituto de Salud Carlos III (AC23_2/00034) and VDR-H by the Agencia Estatal de Investigación (PID2019-109405R and PI-0014-2022). AF-M, IG-R, MLM-C, JCL and JJM were supported by their affiliations. BC-F received unrestricted research funding from Instituto de Salud Carlos III, MINECO, Gobierno de Cantabria, Spanish Network for Research in Mental Health (CIBERSAM), from the Seventh European Union Framework Program and Lundbeck. He has also received honoraria for his participation as a consultant and/or as a speaker at educational events from Janssen Johnson and Johnson, Mylan, Lundbeck, and Otsuka Pharmaceuticals. ACS-H received funding from CIBERSAM (G26) and from the Consejería de Salud y Familias (RH-0063).

## Authors contributions

CM-C contributed to investigation, conceptualization, writing of the original draft, and writing, reviewing, and editing of the manuscript. VDR-H was responsible for investigation, formal analysis, and methodology. AF-M contributed to formal analysis, methodology, and writing of the original draft, as well as writing, reviewing, and editing of the manuscript. IG-R and MLM-C participated in formal analysis. JCL contributed to conceptualization. JJM was involved in conceptualization and methodology. BC-F contributed to conceptualization and oversaw funding acquisition, project administration, supervision, writing, reviewing, and editing of the manuscript. ACS-H contributed to conceptualization, investigation, supervision, writing of the original draft, and writing, reviewing, and editing of the manuscript. All authors have read and approved the final version of the manuscript.

## Abbreviations

THC: Δ9-tetrahydrocannabinol
AD: Alzheimer’s Disease
ApoER2: apolipoprotein E receptor 2
Dab1: disabled homolog-1
CBR: endocannabinoid receptors
ECM: extracellular matrix
GD: gestational day
HP: hippocampus
IPA: Ingenuity Pathway Analysis
MIA: maternal immune activation
Poly(I:C): polyinosinic-polycytidylic acid
PFC: prefrontal cortex
PD: postnatal day
SCZ: schizophrenia
VLDLR: very low-density lipoprotein receptor

## References

Abush, H., Ghose, S., Van Enkevort, E. A., Clementz, B. A., Pearlson, G. D., Sweeney, J. A., Keshavan, M. S., Tamminga, C. A., & Ivleva, E. I. (2018). Associations between adolescent cannabis use and brain structure in psychosis. Psychiatry Research: Neuroimaging, 276, 53–64. 10.1016/j.pscychresns.2018.03.008

Aleman, A., Kahn, R. S., & Selten, J.-P. (2003). Sex Differences in the Risk of Schizophrenia: Evidence From Meta-analysis. Archives of General Psychiatry, 60(6), 565. 10.1001/archpsyc.60.6.565

Alexander, A., Herz, J., & Calvier, L. (2023). Reelin through the years: From brain development to inflammation. Cell Reports, 42(6), 112669. 10.1016/j.celrep.2023.112669

American Psychiatric Association. (2021). Practice guideline for the treatment of patients with Schizophrenia. Third edition. American Psychiatric Publishing.

Ayesa-Arriola, R., De La Foz, V. O.-G., Setién-Suero, E., Ramírez-Bonilla, M. L., Suárez-Pinilla, P., Son, J. M., Vázquez-Bourgon, J., Juncal-Ruiz, M., Gómez-Revuelta, M., Tordesillas-Gutiérrez, D., & Crespo-Facorro, B. (2020). Understanding sex differences in long-term outcomes after a first episode of psychosis. Npj Schizophrenia, 6(1), 33. 10.1038/s41537-020-00120-5

Baines, K. J., Hillier, D. M., Haddad, F. L., Rajakumar, N., Schmid, S., & Renaud, S. J. (2020). Maternal Immune Activation Alters Fetal Brain Development and Enhances Proliferation of Neural Precursor Cells in Rats. Frontiers in Immunology, 11, 1145. 10.3389/fimmu.2020.01145

Bauman, M.D., Van der Water, J. (2020). Translational opportunities in the prenatal immune environment: Promises and limitations of the maternal immune activation model. Neurobiology of Disease, 141, 104864. 10.1016/j.nbd.2020.104864

Bayer, T.A., Falkai, P., Maier, W. (1999). Genetic and non-genetic vulnerability factors in schizophrenia: the basis of the "Two hit hypothesis". Journal of Psychiatric Research, 33, 543–548. 10.1016/s0022-3956(99)00039-4

Bosch, C., Masachs, N., Exposito-Alonso, D., Martínez, A., Teixeira, C. M., Fernaud, I., Pujadas, L., Ulloa, F., Comella, J. X., DeFelipe, J., Merchán-Pérez, A., & Soriano, E. (2016). Reelin Regulates the Maturation of Dendritic Spines, Synaptogenesis and Glial Ensheathment of Newborn Granule Cells. Cerebral Cortex, 26(11), 4282–4298. 10.1093/cercor/bhw216

Carlsson, A., & Carlsson, M. L. (2006). A dopaminergic deficit hypothesis of schizophrenia: The path to discovery. Dialogues in Clinical Neuroscience, 8(1), 137–142. 10.31887/DCNS.2006.8.1/acarlsson

Chesworth, R., & Karl, T. (2017). Molecular Basis of Cannabis-Induced Schizophrenia-Relevant Behaviours: Insights from Animal Models. Current Behavioral Neuroscience Reports, 4(3), 254–279. 10.1007/s40473-017-0120-y

Crespo-Facorro, B., Pelayo-Terán, J. M., Pérez-Iglesias, R., Ramírez-Bonilla, M., Martínez-García, O., Pardo-García, G., & Vázquez-Barquero, J. L. (2007). Predictors of acute treatment response in patients with a first episode of non-affective psychosis: Sociodemographics, premorbid and clinical variables. Journal of Psychiatric Research, 41(8), 659–666. 10.1016/j.jpsychires.2006.05.002

Crespo-Facorro, B., Such, P., Nylander, A.-G., Madera, J., Resemann, H. K., Worthington, E., O’Connor, M., Drane, E., Steeves, S., & Newton, R. (2021). The burden of disease in early schizophrenia – a systematic literature review. Current Medical Research and Opinion, 37(1), 109–121. 10.1080/03007995.2020.1841618

Eranti, S. V., MacCabe, J. H., Bundy, H., & Murray, R. M. (2013). Gender difference in age at onset of schizophrenia: A meta-analysis. Psychological Medicine, 43(1), 155–167. 10.1017/S003329171200089X

Estes, M. L., & McAllister, A. K. (2016). Maternal immune activation: Implications for neuropsychiatric disorders. Science, 353(6301), 772–777. 10.1126/science.aag3194

Fatemi, S. H., Emamian, E. S., Sidwell, R. W., Kist, D. A., Stary, J. M., Earle, J. A., & Thuras, P. (2002). Human influenza viral infection in utero alters glial fibrillary acidic protein immunoreactivity in the developing brains of neonatal mice. Molecular Psychiatry, 7(6), 633–640. 10.1038/sj.mp.4001046

Garey, L. (2010). When cortical development goes wrong: Schizophrenia as a neurodevelopmental disease of microcircuits. Journal of Anatomy, 217(4), 324–333. 10.1111/j.1469-7580.2010.01231.x

Gillespie, B., Panthi, S., Sundram, S., & Hill, R. A. (2024). The impact of maternal immune activation on GABAergic interneuron development: A systematic review of rodent studies and their translational implications. Neuroscience & Biobehavioral Reviews, 156, 105488. 10.1016/j.neubiorev.2023.105488

Glaser, E. M., & Van Der Loos, H. (1981). Analysis of thick brain sections by obverse—Reverse computer microscopy: Application of a new, high clarity Golgi—Nissl stain. Journal of Neuroscience Methods, 4(2), 117–125. 10.1016/0165-0270(81)90045-5

Gray, A., Tattoli, R., Dunn, A., Hodgson, D. M., Michie, P. T., & Harms, L. (2019). Maternal immune activation in mid-late gestation alters amphetamine sensitivity and object recognition, but not other schizophrenia-related behaviours in adult rats. Behavioural Brain Research, 356, 358–364. 10.1016/j.bbr.2018.08.016

Guerrin, C. G. J., Doorduin, J., Sommer, I. E., & De Vries, E. F. J. (2021). The dual hit hypothesis of schizophrenia: Evidence from animal models. Neuroscience & Biobehavioral Reviews, 131, 1150–1168. 10.1016/j.neubiorev.2021.10.025

Guma, E., Plitman, E., Chakravarty, M.M. (2019). The role of maternal immune activation in altering the neurodevelopmental trajectories of offsprings: A translational review of neuroimaging studies with implications for autism spectrum disorders and schizophrenia. Neuroscience & Biobehavioral reviews, 104, 141–157. 10.1016/j.neubiorev.2019.06.020

Guma, E., Cupo, L., Ma, W., Gallino, D., Moquin, L., Gratton, A., Devenyi, G. A., & Chakravarty, M. M. (2023). Investigating the “two-hit hypothesis”: Effects of prenatal maternal immune activation and adolescent cannabis use on neurodevelopment in mice. Progress in Neuro-Psychopharmacology and Biological Psychiatry, 120, 110642. 10.1016/j.pnpbp.2022.110642

Gumusoglu, S.B., Stevens, H.E. (2019). Maternal inflammation and neurodevelopment programming: A review of preclinical ouctomes and implications for translational psychiatry. Biological psychiatry, 15, 85(2);107-121. 10.1016/j.biopsych.2018.08.008

Haddad, F. L., Patel, S. V., & Schmid, S. (2020). Maternal Immune Activation by Poly I:C as a preclinical Model for Neurodevelopmental Disorders: A focus on Autism and Schizophrenia. Neuroscience & Biobehavioral Reviews, 113, 546–567. 10.1016/j.neubiorev.2020.04.012

Haddad, P. M., & Correll, C. U. (2018). The acute efficacy of antipsychotics in schizophrenia: A review of recent meta-analyses. Therapeutic Advances in Psychopharmacology, 8(11), 303–318. 10.1177/2045125318781475

Hall, M. B., Willis, D. E., Rodriguez, E. L., & Schwarz, J. M. (2023). Maternal immune activation as an epidemiological risk factor for neurodevelopmental disorders: Considerations of timing, severity, individual differences, and sex in human and rodent studies. Frontiers in Neuroscience, 17, 1135559. 10.3389/fnins.2023.1135559

Han, V.X., Patel, S., Jones, H.F., Nielsen, T.C., Mohammad, S.S., Hofer, M.J., Gold, W., Brilot, F., Lain, S.J., Nassar, N., Dale, R.C. (2021a). Maternal acute and chronic inflammation in pregnancy is associated with common neurodevelopmental disorders: A systematic review. Translational Psychiatry, 21, 11(1):71. 10.1038/s41398-021-01198-w

Han, V.X., Patel, S., Jones, H.F., Dale, R.C. (2021b). Maternal immune activation and neuroinflammation in human neurodevelopmental disorders. Nature reviews Neurology, 17, 564–579. 10.1038/s41582-021-00530-8

Harte, L. C., & Dow-Edwards, D. (2010). Sexually dimorphic alterations in locomotion and reversal learning after adolescent tetrahydrocannabinol exposure in the rat. Neurotoxicology and Teratology, 32(5), 515–524. 10.1016/j.ntt.2010.05.001

Hattori, M., & Kohno, T. (2021). Regulation of Reelin functions by specific proteolytic processing in the brain. The Journal of Biochemistry, 169(5), 511–516. 10.1093/jb/mvab015

Hitchcock, L. N., Tracy, B. L., Bryan, A. D., Hutchison, K. E., & Bidwell, L. C. (2021). Acute Effects of Cannabis Concentrate on Motor Control and Speed: Smartphone-Based Mobile Assessment. Frontiers in Psychiatry, 11, 623672. 10.3389/fpsyt.2020.623672

Howell, K. R., & Pillai, A. (2016). Long-Term Effects of Prenatal Hypoxia on Schizophrenia-Like Phenotype in Heterozygous Reeler Mice. Molecular Neurobiology, 53(5), 3267–3276. 10.1007/s12035-015-9265-4

Johnson-Ferguson, L., & Di Forti, M. (2023). From heavy cannabis use to psychosis: Is it time to take action? Irish Journal of Psychological Medicine, 40(1), 13–18. 10.1017/ipm.2021.33

Jossin, Y. (2020). Reelin Functions, Mechanisms of Action and Signaling Pathways During Brain Development and Maturation. Biomolecules, 10(6), 964. 10.3390/biom10060964

Jossin, Y., Gui, L., & Goffinet, A. M. (2007). Processing of Reelin by Embryonic Neurons Is Important for Function in Tissue But Not in Dissociated Cultured Neurons. The Journal of Neuroscience, 27(16), 4243–4252. 10.1523/JNEUROSCI.0023-07.2007

Jossin, Y., Ignatova, N., Hiesberger, T., Herz, J., Lambert De Rouvroit, C., & Goffinet, A. M. (2004). The Central Fragment of Reelin, Generated by Proteolytic Processing *In Vivo*, Is Critical to Its Function during Cortical Plate Development. The Journal of Neuroscience, 24(2), 514–521. 10.1523/JNEUROSCI.3408-03.2004

Karamihalev, S., Brivio, E., Flachskamm, C., Stoffel, R., Schimidt, M.V., Chen, A. (2020). Social dominance mediates behavioral adaptation to chronic stress in a sex-specific manner. eLife, 9, e:58723. 10.7554/eLife.58723

Kasten, C. R., Zhang, Y., & Boehm, S. L. (2019). Acute Cannabinoids Produce Robust Anxiety-Like and Locomotor Effects in Mice, but Long-Term Consequences Are Age- and Sex-Dependent. Frontiers in Behavioral Neuroscience, 13, 32. 10.3389/fnbeh.2019.00032

Kirschmann, E. K., Pollock, M. W., Nagarajan, V., & Torregrossa, M. M. (2017). Effects of Adolescent Cannabinoid Self-Administration in Rats on Addiction-Related Behaviors and Working Memory. Neuropsychopharmacology, 42(5), 989–1000. 10.1038/npp.2016.178

Lamanna-Rama N, Romero-Miguel D, Casquero-Veiga M, MacDowell KS, Santa-Marta C, Torres-Sánchez S, Berrocoso E, Leza JC, Desco M, Soto-Montenegro ML. (2024). THC improves behavioural schizophrenia-like deficits that CBD fails to overcome: a comprehensive multilevel approach using the Poly I:C maternal immune activation. Psychiatry Research, 331:115643. doi: 10.1016/j.psychres.2023.115643

Lan, X.-Y., Gu, Y.-Y., Li, M.-J., Song, T.-J., Zhai, F.-J., Zhang, Y., Zhan, J.-S., Böckers, T. M., Yue, X.-N., Wang, J.-N., Yuan, S., Jin, M.-Y., Xie, Y.-F., Dang, W.-W., Hong, H.-H., Guo, Z.-R., Wang, X.-W., & Zhang, R. (2023). Poly(I:C)-induced maternal immune activation causes elevated self-grooming in male rat offspring: Involvement of abnormal postpartum static nursing in dam. Frontiers in Cell and Developmental Biology, 11, 1054381. 10.3389/fcell.2023.1054381

Lecca, S., Luchicchi, A., Scherma, M., Fadda, P., Muntoni, A. L., & Pistis, M. (2019). Δ9-Tetrahydrocannabinol During Adolescence Attenuates Disruption of Dopamine Function Induced in Rats by Maternal Immune Activation. Frontiers in Behavioral Neuroscience, 13, 202. 10.3389/fnbeh.2019.00202

Lidón, L., Urrea, L., Llorens, F., Gil, V., Alvarez, I., Diez-Fairen, M., Aguilar, M., Pastor, P., Zerr, I., Alcolea, D., Lleó, A., Vidal, E., Gavín, R., Ferrer, I., & Del Rio, J. A. (2020). Disease-Specific Changes in Reelin Protein and mRNA in Neurodegenerative Diseases. Cells, 9(5), 1252. 10.3390/cells9051252

MacDowell KS, Munarriz-Cuezva E, Caso JR, Madrigal JL, Zabala A, Meana JJ, García-Bueno B, Leza JC. (2017). Paliperidone reverts Toll-like receptor 3 signaling pathway activation and cognitive deficits in a maternal immune activation mouse model of schizophrenia. Neuropharmacology, 116:196–207. doi: 10.1016/j.neuropharm.2016.12.025

Malaspina, D., Sohler, N. L., Susser, E. S. (1999). Interaction of genes and prenatal exposures in schizophrenia. Prenatal exposures in schizophrenia, American Psychiatric Association, (pp. 35–59).

Manza, P., Yuan, K., Shokri-Kojori, E., Tomasi, D., & Volkow, N. D. (2020). Brain structural changes in cannabis dependence: Association with MAGL. Molecular Psychiatry, 25(12), 3256–3266. 10.1038/s41380-019-0577-z

Martín-Cuevas, C., Ramos-Herrero, V. D., Crespo-Facorro, B., & Sánchez-Hidalgo, A. C. (2023). Prenatal risk factors and postnatal cannabis exposure: Assessing dual models of schizophrenia-like rodents. Neuroscience & Biobehavioral Reviews, 154, 105409. 10.1016/j.neubiorev.2023.105409

Marzan, S., Aziz, Md. A., & Islam, M. S. (2021). Association Between REELIN Gene Polymorphisms (rs7341475 and rs262355) and Risk of Schizophrenia: An Updated Meta-analysis. Journal of Molecular Neuroscience, 71(4), 675–690. 10.1007/s12031-020-01696-4

Mashhoon, Y., Sava, S., Sneider, J. T., Nickerson, L. D., & Silveri, M. M. (2015). Cortical thinness and volume differences associated with marijuana abuse in emerging adults. Drug and Alcohol Dependence, 155, 275–283. 10.1016/j.drugalcdep.2015.06.016

Maynard, T.M., Sikich, L., Lieberman, J.A., LaMantia, A. S. (2001). Neural Development, Cell-Cell Signaling, and the “Two-Hit” Hypothesis of Schizophrenia. Schizophrenia Bulletin, 27, 457–476, 10.1093/oxfordjournals.schbul.a006887

McCutcheon, R. A., Reis Marques, T., & Howes, O. D. (2020). Schizophrenia—An Overview. JAMA Psychiatry, 77(2), 201. 10.1001/jamapsychiatry.2019.3360

Mendrek, A., & Mancini-Marïe, A. (2016). Sex/gender differences in the brain and cognition in schizophrenia. Neuroscience & Biobehavioral Reviews, 67, 57–78. 10.1016/j.neubiorev.2015.10.013

Meyer, U., Nyffeler, M., Engler, A., Urwyler, A., Schedlowski, M., Knuesel, I., Yee, B. K., & Feldon, J. (2006). The Time of Prenatal Immune Challenge Determines the Specificity of Inflammation-Mediated Brain and Behavioral Pathology. The Journal of Neuroscience, 26(18), 4752–4762. 10.1523/JNEUROSCI.0099-06.2006

Moreno-Sancho, L., Juncal-Ruiz, M., Vázquez-Bourgon, J., Ortiz-Garcia De La Foz, V., Mayoral-van Son, J., Tordesillas-Gutierrez, D., Setien-Suero, E., Ayesa-Arriola, R., & Crespo-Facorro, B. (2022). Naturalistic study on the use of clozapine in the early phases of non-affective psychosis: A 10-year follow-up study in the PAFIP-10 cohort. Journal of Psychiatric Research, 153, 292–299. 10.1016/j.jpsychires.2022.07.015

Morrens, M., Hulstijn, W., Lewi, P. J., De Hert, M., & Sabbe, B. G. C. (2006). Stereotypy in schizophrenia. Schizophrenia Research, 84(2-3), 397–404. 10.1016/j.schres.2006.01.024

Mueller, F. S., Richetto, J., Hayes, L. N., Zambon, A., Pollak, D. D., Sawa, A., Meyer, U., & Weber-Stadlbauer, U. (2019). Influence of poly(I:C) variability on thermoregulation, immune responses and pregnancy outcomes in mouse models of maternal immune activation. Brain, Behavior, and Immunity, 80, 406–418. 10.1016/j.bbi.2019.04.019

Munarriz-Cuezva E, Meana JJ. (2025). Poly (I:C)-induced maternal immune activation generates impairment of reversal learning performance in offspring. Journal of Neurochemistry,169(1):e16212. doi: 10.1111/jnc.16212

Murlanova, K., & Pletnikov, M. V. (2023). Modeling psychotic disorders: Environment x environment interaction. Neuroscience & Biobehavioral Reviews, 152, 105310. 10.1016/j.neubiorev.2023.105310

Nenadic, I., Yotter, R. A., Sauer, H., & Gaser, C. (2015). Patterns of cortical thinning in different subgroups of schizophrenia. British Journal of Psychiatry, 206(6), 479–483. 10.1192/bjp.bp.114.148510

Ogino, H., Hisanaga, A., Kohno, T., Kondo, Y., Okumura, K., Kamei, T., Sato, T., Asahara, H., Tsuiji, H., Fukata, M., & Hattori, M. (2017). Secreted Metalloproteinase ADAMTS-3 Inactivates Reelin. The Journal of Neuroscience, 37(12), 3181–3191. 10.1523/JNEUROSCI.3632-16.2017

Okugawa, E., Ogino, H., Shigenobu, T., Yamakage, Y., Tsuiji, H., Oishi, H., Kohno, T., & Hattori, M. (2020). Physiological significance of proteolytic processing of Reelin revealed by cleavage-resistant Reelin knock-in mice. Scientific Reports, 10(1), 4471. 10.1038/s41598-020-61380-w

Ovadia, G., & Shifman, S. (2011). The Genetic Variation of RELN Expression in Schizophrenia and Bipolar Disorder. PLoS ONE, 6(5), e19955. 10.1371/journal.pone.0019955

Patel, K. R., Cherian, J., Gohil, K., & Atkinson, D. (2014). Schizophrenia: Overview and Treatment Options.

Patel, S. J., Khan, S., M, S., & Hamid, P. (2020). The Association Between Cannabis Use and Schizophrenia: Causative or Curative? A Systematic Review. Cureus. 10.7759/cureus.9309

Prashad, S., & Filbey, F. M. (2017). Cognitive motor deficits in cannabis users. Current Opinion in Behavioral Sciences, 13, 1–7. 10.1016/j.cobeha.2016.07.001

Rais, M., Van Haren, N. E. M., Cahn, W., Schnack, H. G., Lepage, C., Collins, L., Evans, A. C., Hulshoff Pol, H. E., & Kahn, R. S. (2010). Cannabis use and progressive cortical thickness loss in areas rich in CB1 receptors during the first five years of schizophrenia. European Neuropsychopharmacology, 20(12), 855–865. 10.1016/j.euroneuro.2010.08.008

Ratnayake, U., Quinn, T. A., Castillo-Melendez, M., Dickinson, H., & Walker, D. W. (2012). Behaviour and hippocampus-specific changes in spiny mouse neonates after treatment of the mother with the viral-mimetic Poly I:C at mid-pregnancy. *Brain*, Behavior, and Immunity, 26(8), 1288–1299. 10.1016/j.bbi.2012.08.011

Reisinger, S., Khan, D., Kong, E., Berger, A., Pollak, A., & Pollak, D. D. (2015). The Poly(I:C)-induced maternal immune activation model in preclinical neuropsychiatric drug discovery. Pharmacology & Therapeutics, 149, 213–226. 10.1016/j.pharmthera.2015.01.001

Rodríguez-Sánchez, J. M., Crespo-Facorro, B., González-Blanch, C., Pérez-Iglesias, R., Vázquez-Barquero, J. L., & PAFIP Group Study. (2007). Cognitive dysfunction in first-episode psychosis: The processing speed hypothesis. British Journal of Psychiatry, 191(S51), s107–s110. 10.1192/bjp.191.51.s107

Rogers, D. C., Fisher, E. M. C., Brown, S. D. M., Peters, J., Hunter, A. J., & Martin, J. E. (1997). Behavioral and functional analysis of mouse phenotype: SHIRPA, a proposed protocol for comprehensive phenotype assessment. Mammalian Genome, 8(10), 711–713. 10.1007/s003359900551

Sánchez-Hidalgo, A. C., Martín-Cuevas, C., Crespo-Facorro, B., & Garrido-Torres, N. (2022). Reelin Alterations, Behavioral Phenotypes, and Brain Anomalies in Schizophrenia: A Systematic Review of Insights From Rodent Models. Frontiers in Neuroanatomy, 16, 844737. 10.3389/fnana.2022.844737

Schwartzer, J. J., Careaga, M., Onore, C. E., Rushakoff, J. A., Berman, R. F., & Ashwood, P. (2013). Maternal immune activation and strain specific interactions in the development of autism-like behaviors in mice. Translational Psychiatry, 3(3), e240–e240. 10.1038/tp.2013.16

Setién-Suero, E., Suárez-Pinilla, P., Ferro, A., Tabarés-Seisdedos, R., Crespo-Facorro, B., & Ayesa-Arriola, R. (2020). Childhood trauma and substance use underlying psychosis: A systematic review. European Journal of Psychotraumatology, 11(1), 1748342. 10.1080/20008198.2020.1748342

Shangase, K. B., Luvuno, M., & Mabandla, M. V. (2023). Investigating the Robustness of a Rodent “Double Hit” (Post-Weaning Social Isolation and NMDA Receptor Antagonist) Model as an Animal Model for Schizophrenia: A Systematic Review. Brain Sciences, 13(6), 848. 10.3390/brainsci13060848

Smith, S. E. P., Elliott, R. M., & Anderson, M. P. (2012). Maternal Immune Activation Increases Neonatal Mouse Cortex Thickness and Cell Density. Journal of Neuroimmune Pharmacology, 7(3), 529–532. 10.1007/s11481-012-9372-1

Smith-Osbone, L., Duong, A., Resendez, A., Palme, R., Fadok, J.P. (2023). Female dominance hierarchies influence responses to psychosocial stressors. Current Biology, 33, 1535–1549. 10.1016/j.cub.2023.03.020

Suárez-Pinilla, P., López-Gil, J., & Crespo-Facorro, B. (2014). Immune system: A possible nexus between cannabinoids and psychosis. Brain, Behavior, and Immunity, 40, 269–282. 10.1016/j.bbi.2014.01.018

Valderrama-Mantilla, A. I., Martín-Cuevas, C., Gómez-Garrido, A., Morente-Montilla, C., Crespo-Facorro, B., & García-Cerro, S. (2025). Shared molecular signature in Alzheimer’s disease and schizophrenia: A systematic review of the reelin signaling pathway. Neuroscience & Biobehavioral Reviews, 169, 106032. 10.1016/j.neubiorev.2025.106032

Van der Berg, W.E., Lamballais, S., Kushner, S.A. (2015). Sex-specific mechanism of social hierarchy in mice. Neuropsychopharmacology, 40, 1364–1372. 10.1038/npp.2014.319

Vlasova RM, Iosif AM, Ryan AM, Funk LH, Murai T, Chen S, Lesh TA, Rowland DJ, Bennett J, Hogrefe CE, Maddock RJ, Gandal MJ, Geschwind DH, Schumann CM, Van de Water J, McAllister AK, Carter CS, Styner MA, Amaral DG, Bauman MD. (2021). Maternal immune activation during pregnancy alters postnatal brain growth and cognitive development in nonhuman primate offspring. Journal of Neuroscience, 1;41(48):9971-9987. 10.1523/JNEUROSCI.0378-21.2021

Wasser, C. R., & Herz, J. (2017). Reelin: Neurodevelopmental Architect and Homeostatic Regulator of Excitatory Synapses. Journal of Biological Chemistry, 292(4), 1330–1338. 10.1074/jbc.R116.766782

Williamson, C.M., Lee, W., DeCasien, A.R., Lanham, A., Romeo, R.D., Curley, J.P. (2019). Social hierarchy position in female mice is associated with plasma corticosterone levels and hypothalamic gene expression. Scientific reports, 13, 7324. 10.1038/s41598-019-43747-w

Yamakage, Y., Kato, M., Hongo, A., Ogino, H., Ishii, K., Ishizuka, T., Kamei, T., Tsuiji, H., Miyamoto, T., Oishi, H., Kohno, T., & Hattori, M. (2019). A disintegrin and metalloproteinase with thrombospondin motifs 2 cleaves and inactivates Reelin in the postnatal cerebral cortex and hippocampus, but not in the cerebellum. Molecular and Cellular Neuroscience, 100, 103401. 10.1016/j.mcn.2019.103401

Yan, J., Cui, Y., Li, Q., Tian, L., Liu, B., Jiang, T., Zhang, D., & Yan, H. (2019). Cortical thinning and flattening in schizophrenia and their unaffected parents. *Neuropsychiatric Disease and Treatment*, Volume 15, 935–946. 10.2147/NDT.S195134

