## Supplementary material for "Disruption of Reelin signaling in a dual-hit mouse model of schizophrenia: impact of postnatal Δ9-tetrahydrocannabinol exposure in a maternal immune activation model": Supplementary material 1.docx

1. **Maternal immune activation evaluation**

**1.1 Quality control of the Poly(I:C) effect through assessment of maternal weight and temperature**

Previous studies have reported significant weight loss in pregnant dams 24 hours after Poly(I:C) administration (Chow et al., 2016; Lecca et al., 2019). A two-way ANOVA revealed significant effects of days (F(4,68) = 11.26, p < 0.0001), Poly(I:C) (F(1,17) = 12.35, p < 0.01) and days x Poly(I:C) interaction (F(4,68) = 6.42, p < 0.001). Post hoc comparisons (Sidak’s test) showed that pregnant mice exhibited significantly reduced body weight 24 (p < 0.0001) and 48 (p < 0.001) hours after Poly(I:C) injections compared to controls injected with saline (Fig. S1A).

Secondly, it has been reported that the temperature of females injected with Poly(I:C) decreases by 0.5 ºC to 1 ºC at 3 hours and progressively increases at 6 and 24 hours, finally returning to basal temperature at 48 hours (Chow et al., 2016; Mueller et al., 2019). The statistical analysis showed significant effects of days (F(4,68) = 35.19, p < 0.0001), Poly(I:C) (F(1,17) = 6.14, p < 0.05) and days x Poly(I:C) interaction (F(4,68) = 37.86, p < 0.0001). Poly(I:C)-treated dams exhibited a significant reduction in temperature 3 hours after injection (p < 0.0001) and a significant increase in temperature 24 hours after injection (p < 0.0001) compared to saline-treated controls (Fig. S1B). In summary, MIA induced by Poly(I:C) injection in pregnant dams has been validated through the assessment of their weight and temperature.

**1.2 Cytokines levels are increased in maternal spleen and embryos**

The acute inflammatory response was examined in pregnant dams and embryos (including placenta) using RT-qPCR 3 hours after Poly(I:C) injection. We observed significantly increased levels of IL-6 (p < 0.001) and TNFα (p < 0.001) in the maternal spleen of Poly(I:C)-injected dams compared to saline controls (Fig. S1C) (Student t-test). Similar results were observed in the embryos (IL-6: p < 0.0001; TNFα: p < 0.01) (Fig. S1D), corroborating the inflammatory response. These results are consistent with previous studies (Chow et al., 2016; Hameete et al., 2021), thus confirming the maternal immune response in the animals.

2. **The dual-hit paradigm does not affect body weight**

Cannabis consumption is linked to alterations in appetite and body weight (Williams & Kirkham, 2002; Farrimond et al., 2012; Fearby et al., 2022). However, there were no significant differences in body weight among any treatment groups either prior to THC treatment (PD55) or post-treatment (PD60). At PD75, the three-way ANOVA revealed a significant effect of sex (F(1,76) = 321.8, p < 0.0001) and THC (F(1,76) = 12.85, p < 0.001). Separate two-way ANOVAs were conducted for males and females. In males, a significant effect of THC was found (F(1,38) = 9.68, p < 0.01), and post-hoc analysis showed a decreased body weight in S+T mice compared to P+V mice (p < 0.05), but no effect was observed in the dual-hit group. Two-way ANOVA in female body weight did not report any significant effect (Fig. S1E).

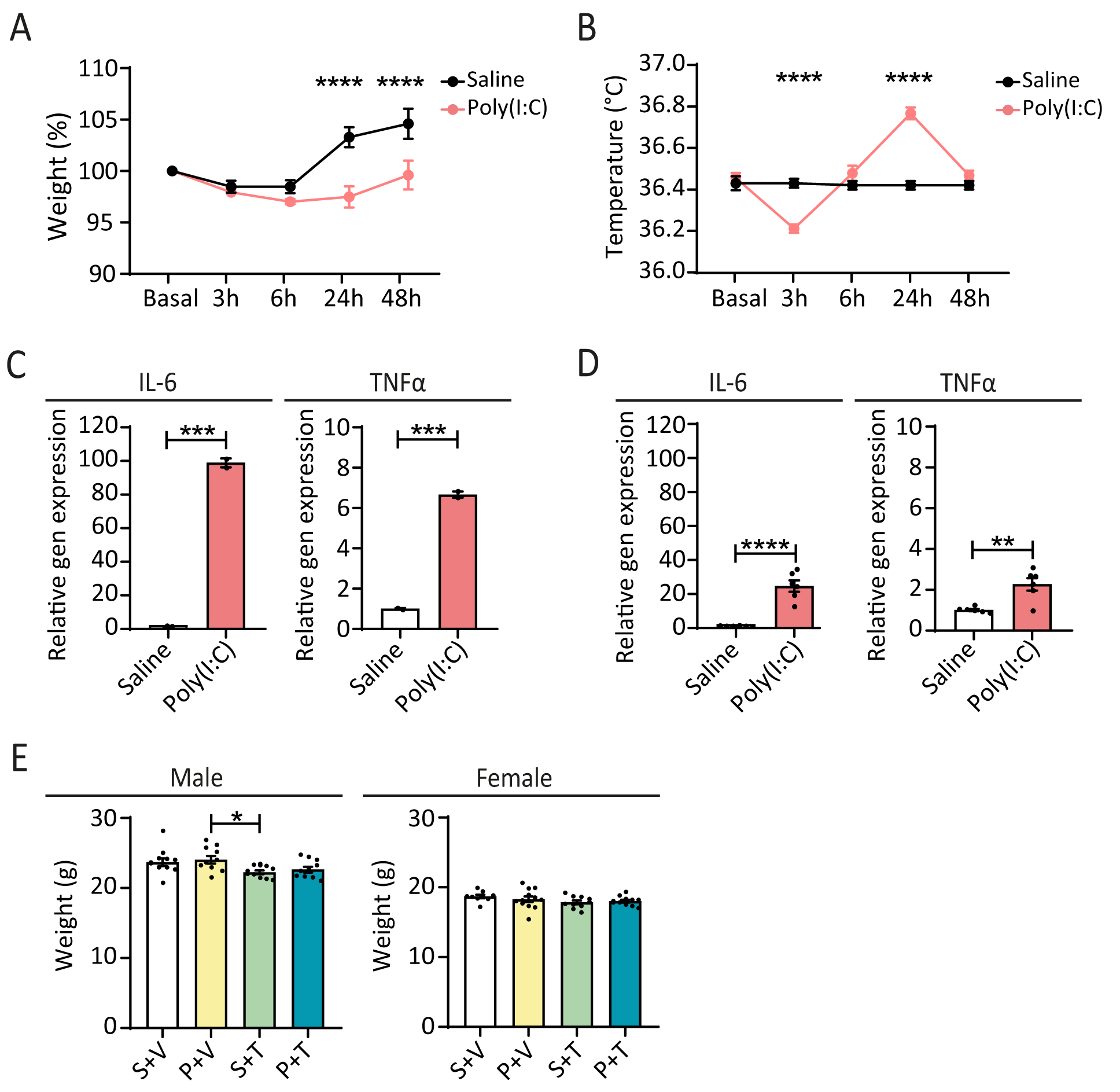

**Figure S1.** MIA evaluation and THC effect on body weight. **A,** Body weight trajectories of pregnant dams before saline or Poly(I:C) injection (basal) and 3, 6 24 and 48-hours post-injection (Saline: n = 10; Poly(I:C): n = 9) (repeated-measures two-way ANOVA with Sidak’s post-hoc analysis; ****p < 0.0001). **B,** Temperature trajectories of pregnant dams before saline or Poly(I:C) injection (basal) and 3, 6, 24 and 48-hours post-injection (Saline: n = 10; Poly(I:C): n = 9) (repeated-measures two-way ANOVA with Sidak’s post-hoc analysis; ****p < 0.0001). **C,** Cytokines levels in maternal spleen (n = 2 per treatment) (Student t-test; IL-6: ***p < 0.001, TNF: ***p < 0.001). **D,** Cytokines levels in embryos with placenta (n = 6 per treatment) (Student t-test; IL-6: ****p < 0.0001, TNF: **p = 0.002). **E,** Adult body weight (n = 9-12 per treatment and sex) (two-way ANOVA with Tukey’s post-hoc analysis; *p < 0.05). Abbreviations: Saline + Vehicle (S+V), Poly(I:C) + Vehicle (P+V), Saline + THC (S+T), Poly(I:C) + THC (P+T).

3. **Behavioral Assays**
   3.1 **Execution order of behavioral tests**

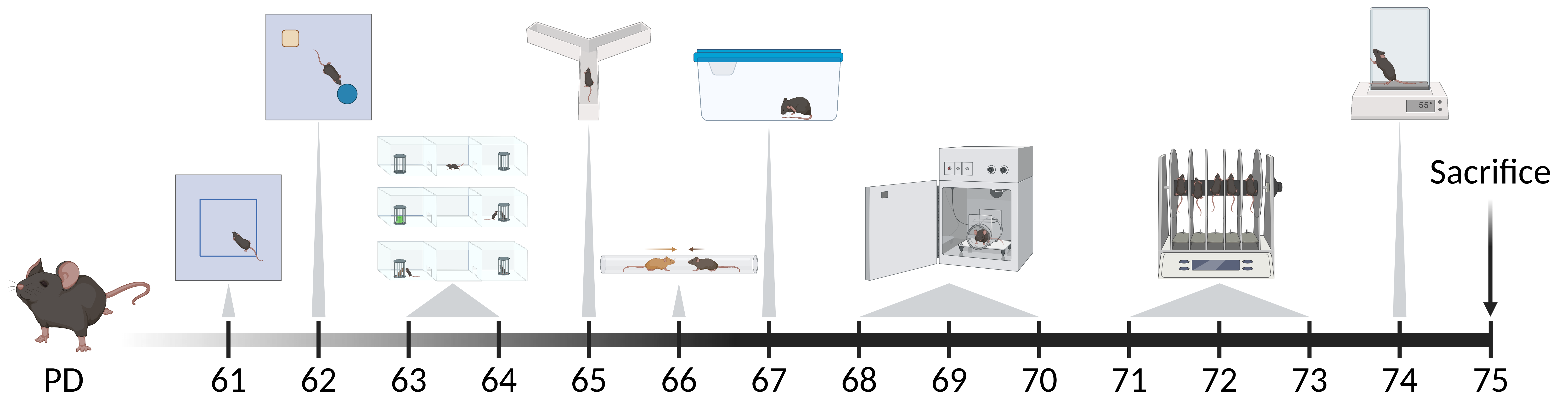

**Figure S2.** Execution order of behavioral tests from the least to most stressful: open field, novel object recognition, 3-chamber test, y-maze, tube test, self-grooming, startle reflex and prepulse inhibition, rotarod and hotplate. Created with Biorender.com.

**3.2 Open field test**

The open field test was used to assess locomotor activity and anxiety-like behavior. Animals were placed in a 50 x 50 cm arena for 15 minutes, and their movement was recorded and analyzed using Viewer software (Biobserve). The central area was defined as a 30 x 30 cm zone, with the remaining area designated as the peripheral zone. Total distance traveled and mean velocity were analyzed.

**3.3 Self-grooming test**

The self-grooming test was performed to assess stereotyped behavior. Mice were habituated in a clean home cage without bedding for 10 minutes, followed by a 10-minutes recording period. Both the grooming time and the number of grooming events were quantified.

**3.4 Startle reflex and prepulse inhibition**

The StartFear Combined system (LE0823G, Panlab Harvard Apparatus) was used to perform the test as previously described with minor modifications (García-Partida et al., 2022). The startle response (SR) was automatically analyzed with the Packwin software (Panlab Harvard Apparatus). Mice were placed in a restraint device and exposed to a constant background white noise of 70 dB. Each session consisted of a 10-minute exploration period, followed by 5 pulse-alone (120 dB). Then, mice were exposed to a pseudorandom block of 50 trials including no stimulus, pulse-alone (120 dB) and 3 combinations of prepulse-pulse trials (74, 80 or 86 dB + 120 dB), with inter-trials intervals of 10, 15, or 20 s. Finally, they were subjected to 25 pulses of different intensities (0, 90, 100, 110 and 120 dB). Pulses lasted 40 ms, and prepulses lasted 20 ms. The percentage of prepulse inhibition (%PPI) was calculated for each prepulse-pulse combination as [(mean SR to pulse-alone - mean SR to prepulse-pulse combination) / mean SR to pulse-alone] x 100.

**3.5 Rotarod**

Motor coordination and learning were evaluated using the rotarod (LE8205, Panlab Harvard Apparatus). On the first day, mice were trained to walk forward on the rod at 4 rpm for 3 minutes. This habituation was performed 3 times per day for 1 minute before each test the session. The test phase comprised three trials per day over three consecutive days. The apparatus was programmed to accelerate from 4 to 40 rpm over 5 minutes, and the latency to fall was automatically recorded using Sedacom software (Panlab Harvard Apparatus).

**3.6 Hotplate**

The hotplate test (LE7406, Panlab Harvard Apparatus) was conducted to assess heat sensitivity using a temperature of 55 ºC. The latency for mice to jump or lick their paws was analyzed.

**3.7 Three-chamber test**

The three-chamber test was used to evaluate social preference and social novelty and was performed as described previously with minor modifications (Silverman et al., 2010). The test chamber was divided into three compartments, each measuring 42 x 20 cm, with a total size of 42 x 60 cm (LE894T, Panlab Harvard Apparatus). The test comprised three 10-minute phases. In the exploration phase, animals were allowed to explore the arena. In the social preference phase, an unfamiliar mouse and an object were placed in the cages. In the social novelty phase, mice interacted with the previously encountered familiar mouse and a new, unfamiliar mouse. The interaction time with the object or familiar mouse or familiar vs new mouse was quantified. Social and memory index were calculated as [(interaction time with mouse or new mouse - interaction time with object or familiar mouse) / (interaction time with mouse or new mouse + interaction time with object or familiar mouse)].

**3.8 Tube test**

The tube test was conducted to evaluate dominance. Pairs of same-sex mice with different treatments were placed at opposite ends of a 2.6-cm diameter plexiglass tube and given 2 minutes to interact. The first animal to exit the tube was recorded as the dominant mouse. Each mouse was tested against two different animals with varying treatments.

**3.9 Novel object recognition test**

Mice were placed in the open field chamber containing two identical objects and allowed to explore for 10 minutes. After a 3-hour delay, one of the objects was replaced with a novel object and mice were given another 10 minutes to explore the arena. The percentage of time spent interacting with the familiar and novel objects was quantified. The memory index was calculated as [(interaction time with new object - interaction time with familiar object) / (interaction time with new object + interaction time with familiar object)].

**3.10 Y-maze test**

Mice were allowed to explore a y-maze, with each arm measuring 6 x 30 cm, for 5 minutes while their movement was recorded. Spontaneous alternation was analyzed using SMART software (Panlab Harvard Apparatus).

4. **Heat sensitivity in the hot plate test**

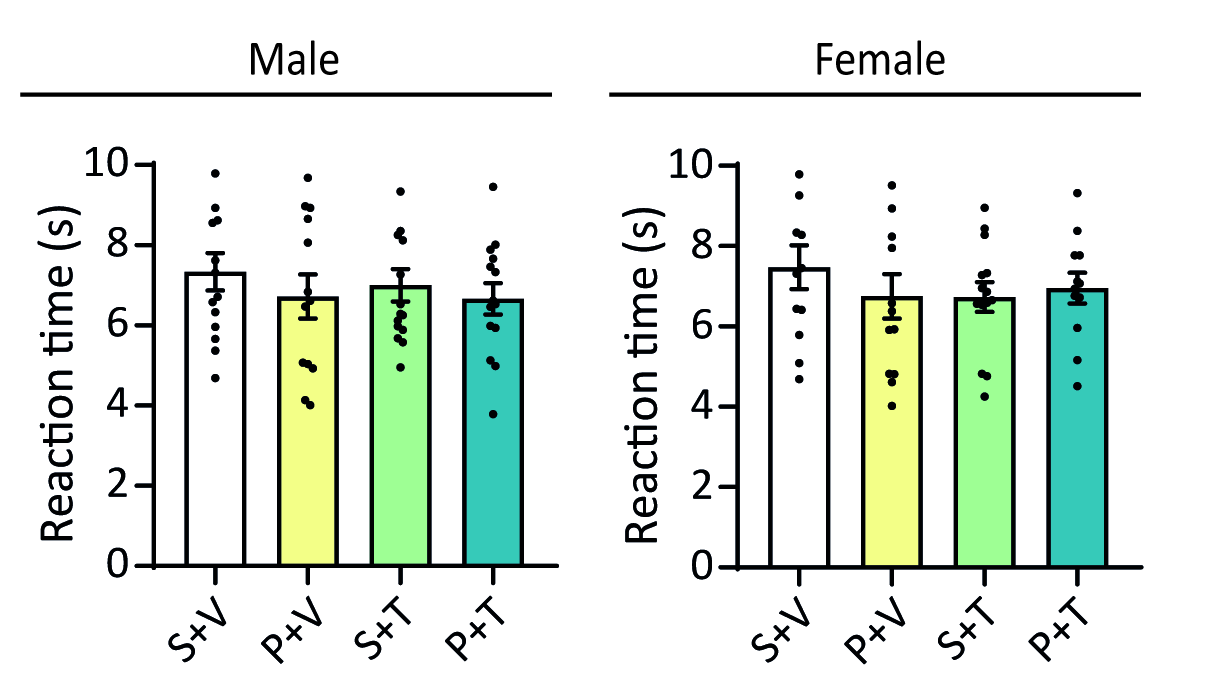

**Figure S3.** Analysis of heat sensitivity in the hot plate test. Reaction time (s) (n = 12-15 per treatment and sex) (three-way ANOVA). Abbreviations: Saline + Vehicle (S+V), Poly(I:C) + Vehicle (P+V), Saline + THC (S+T), Poly(I:C) + THC (P+T).

5. **Startle reflex and prepulse inhibition**

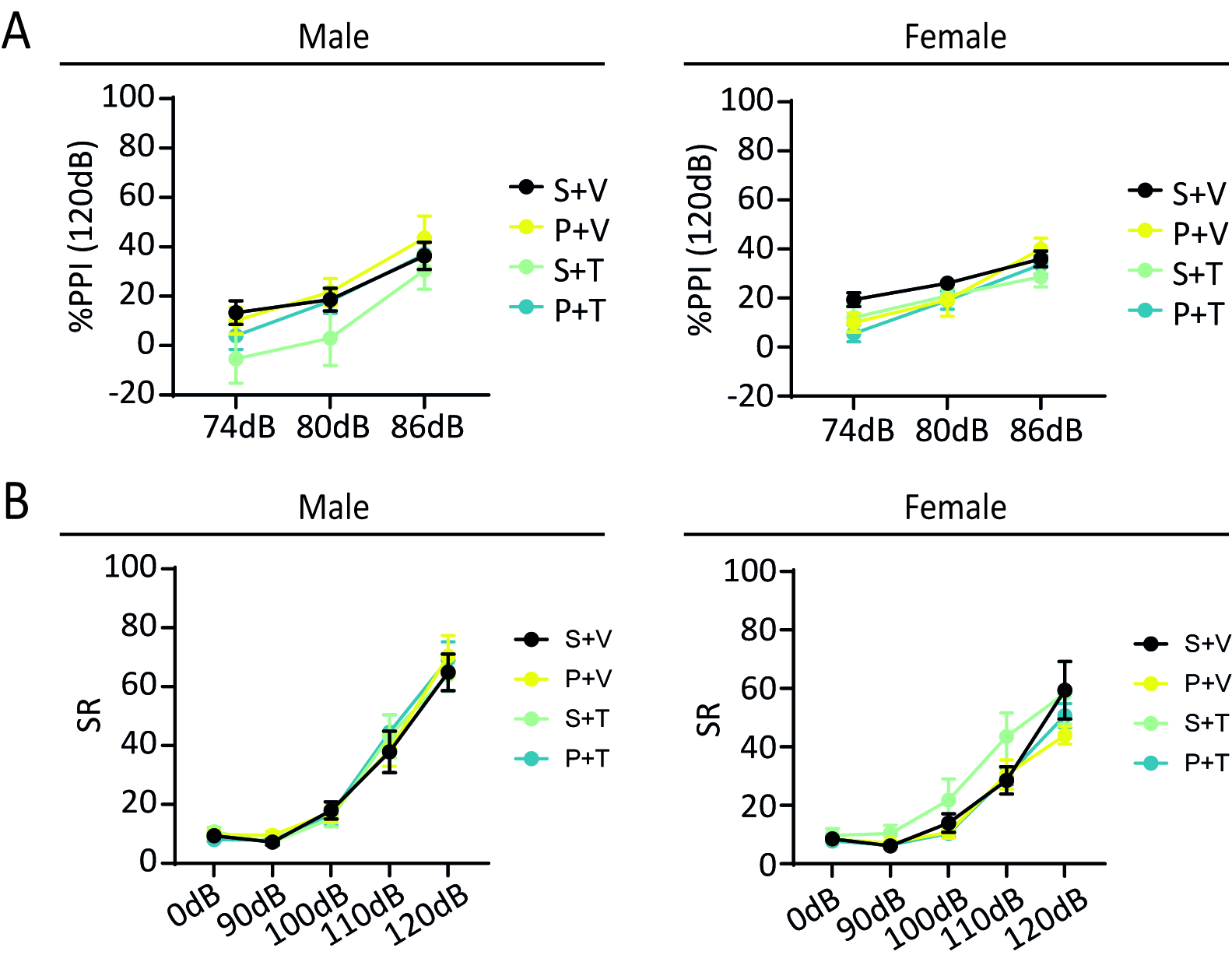

**Figure S4.** Analysis of startle reflex and prepulse inhibition. **A,** Prepulse inhibition (%). **B,** Startle response (SR) (%) (n = 6-11 per treatment and sex) (repeated measures three-way ANOVA with Tukey’s post-hoc analysis). Abbreviations: Saline + Vehicle (S+V), Poly(I:C) + Vehicle (P+V), Saline + THC (S+T), Poly(I:C) + THC (P+T).

6. **Analysis of paired treatments in the tube test**

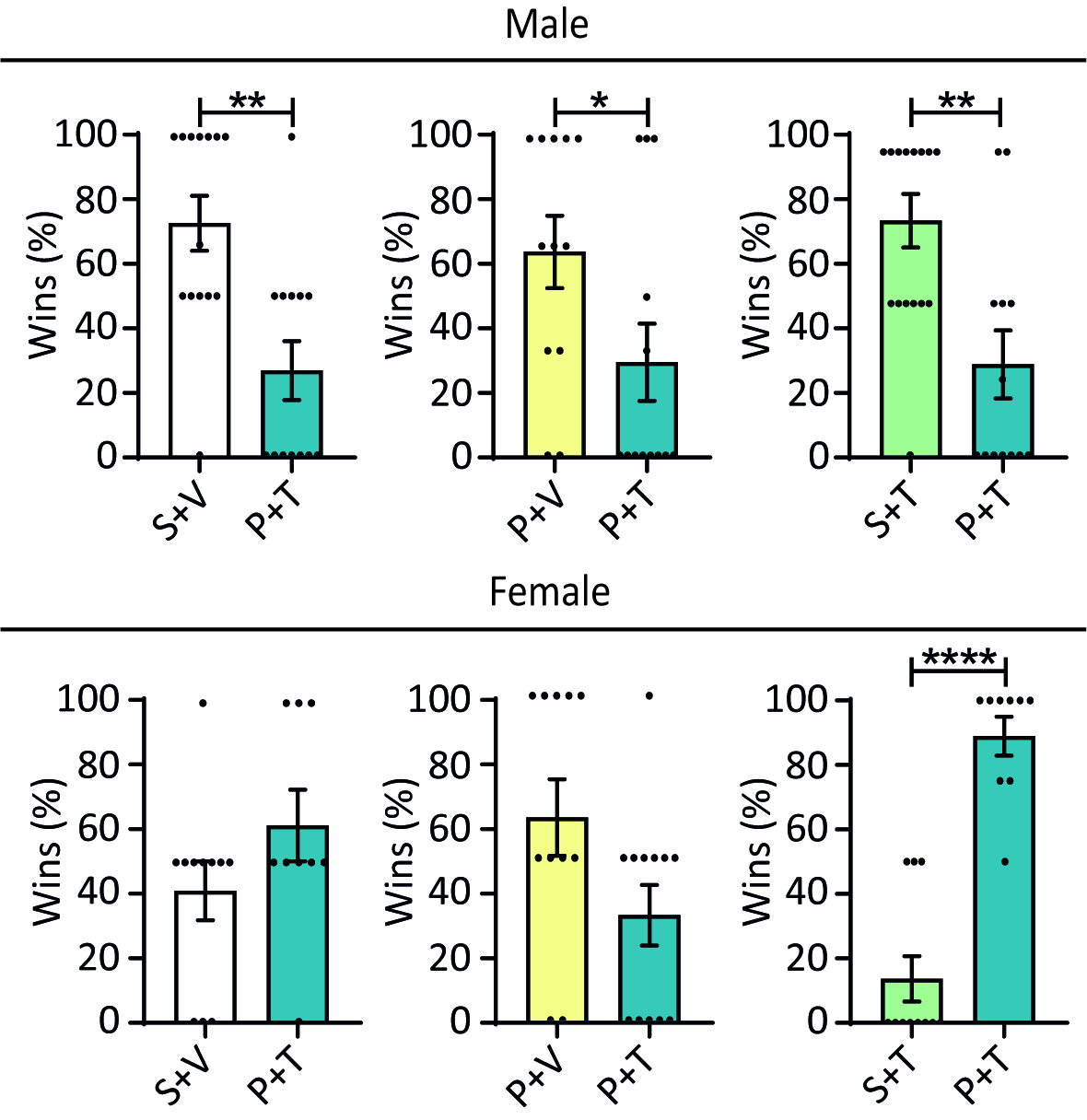

**Figure S5.** Analysis of paired treatment in the tube test. Percentage of wins (n = 12-15 per treatment and sex) (Student t-test; *p < 0.05, **p<0.01, ****p < 0.0001). Abbreviations: Saline + Vehicle (S+V), Poly(I:C) + Vehicle (P+V), Saline + THC (S+T), Poly(I:C) + THC (P+T).

7. **Spontaneous alternation in the y-maze test**

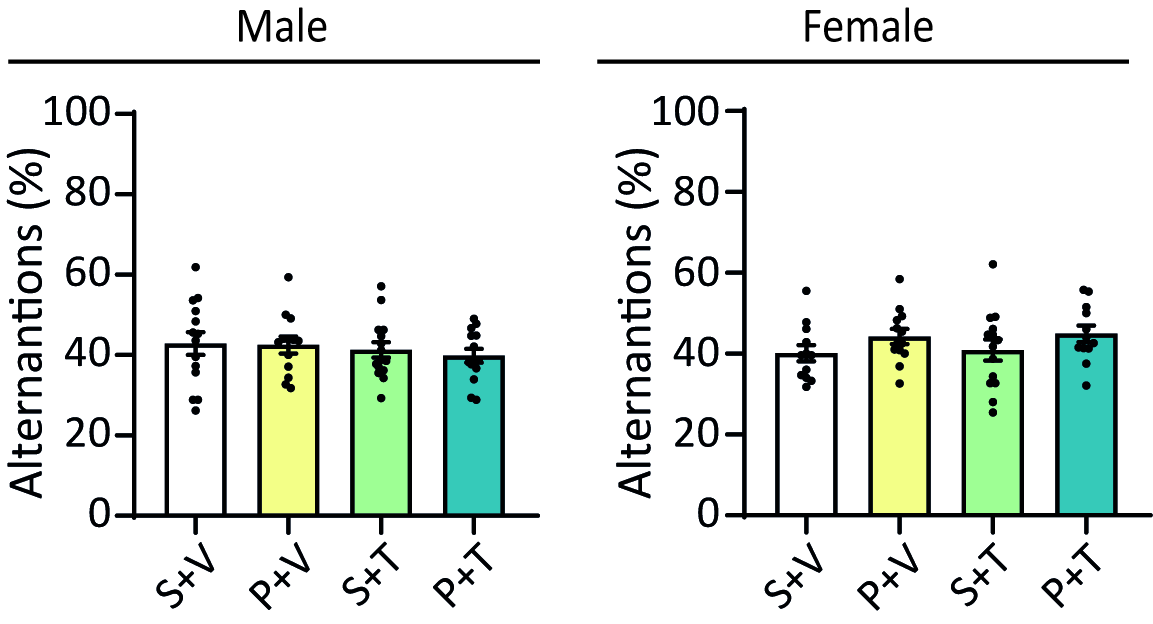

**Figure S6.** Analysis of spontaneous alternations in the y-maze. Percentage of alternations (n = 12-15 per treatment and sex) (three-way ANOVA). Abbreviations: Saline + Vehicle (S+V), Poly(I:C) + Vehicle (P+V), Saline + THC (S+T), Poly(I:C) + THC (P+T).

8. **Proteomic analysis of S+T vs S+V**

**Figure S7. Quantitative proteomics analysis of PFC samples from Saline + THC-treated mice.** **A,** Venn diagram showing the number and percentage of proteins altered after Saline + THC treatment in males (yellow) and females (blue), including those commonly affected in both sexes. **B,** Volcano plot illustrating the distribution of differentially expressed proteins based on log2(Fold Change) and -log10(p-value) for the Saline + THC male vs. Saline + THC female comparison, normalized to their respective controls. Proteins with significant differential expression (p < 0.05) are highlighted in red. **C,** Ingenuity Canonical Pathways enrichment analysis depicting significantly differentially expressed pathways in Saline + THC-treated males (upper panel) and females (bottom panel), normalized to their respective controls. Dot size corresponds to the number of altered proteins within each pathway (n molecules). **D,** Dot plot quantifying the number of significantly altered proteins per pathway set and the total dataset (Gene Ratio) in males (upper panel) and females (bottom panel), normalized to their respective controls. Color intensity represents the adjusted p-value (-log10), while dot size indicates the number of altered proteins within each pathway. Abbreviations: Saline + Vehicle (S+V), Poly(I:C) + Vehicle (P+V), Saline + THC (S+T), Poly(I:C) + THC (P+T).

9. **References**

Chow, K.-H., Yan, Z., & Wu, W.-L. (2016). Induction of Maternal Immune Activation in Mice at Mid-gestation Stage with Viral Mimic Poly(I:C). *Journal of Visualized Experiments*, *109*, 53643. https://doi.org/10.3791/53643

Farrimond, J. A., Whalley, B. J., & Williams, C. M. (2012). Cannabinol and cannabidiol exert opposing effects on rat feeding patterns. *Psychopharmacology*, *223*(1), 117-129. https://doi.org/10.1007/s00213-012-2697-x

Fearby, N., Penman, S., & Thanos, P. (2022). Effects of Δ9-Tetrahydrocannibinol (THC) on Obesity at Different Stages of Life: A Literature Review. *International Journal of Environmental Research and Public Health*, *19*(6), 3174. https://doi.org/10.3390/ijerph19063174

García-Partida, J.A., Torres-Sánchez, S., MacDowell, K., Fernández-Ponce, M.T., Casas, L., Mantell, C., Soto-Montenegro, M.L., Romero-Miguel, D., Lamanna-Rama, N., Leza, J.C., Desco, M., Berrocoso, E. The effects of mando leaf extract during adolescence and adulthood in a rat model of schizophrenia. *Frontiers in Pharmacology, 13*:886514. https://doi.org/10.3389/fphar.2022.886514.

Hameete, B. C., Fernández-Calleja, J. M. S., De Groot, M. W. G. D. M., Oppewal, T. R., Tiemessen, M. M., Hogenkamp, A., De Vries, R. B. M., & Groenink, L. (2021). The poly(I:C)-induced maternal immune activation model; a systematic review and meta-analysis of cytokine levels in the offspring. *Brain, Behavior, & Immunity - Health*, *11*, 100192. https://doi.org/10.1016/j.bbih.2020.100192

Lecca, S., Luchicchi, A., Scherma, M., Fadda, P., Muntoni, A. L., & Pistis, M. (2019). Δ9-Tetrahydrocannabinol During Adolescence Attenuates Disruption of Dopamine Function Induced in Rats by Maternal Immune Activation. *Frontiers in Behavioral Neuroscience*, *13*, 202. https://doi.org/10.3389/fnbeh.2019.00202

Mueller, F. S., Richetto, J., Hayes, L. N., Zambon, A., Pollak, D. D., Sawa, A., Meyer, U., & Weber-Stadlbauer, U. (2019). Influence of poly(I:C) variability on thermoregulation, immune responses and pregnancy outcomes in mouse models of maternal immune activation. *Brain, Behavior, and Immunity*, *80*, 406-418. https://doi.org/10.1016/j.bbi.2019.04.019

Silverman, J. L., Yang, M., Lord, C., & Crawley, J. N. (2010). Behavioural phenotyping assays for mouse models of autism. *Nature Reviews Neuroscience*, *11*(7), 490-502. https://doi.org/10.1038/nrn2851

Williams, C. M., & Kirkham, T. C. (2002). *Observational analysis of feeding induced by D9-THC and anandamide*.
